# A human gut metagenome-assembled genome catalogue spanning 41 countries supports genome-scale metabolic models

**DOI:** 10.1101/2024.12.11.627901

**Authors:** Junyeong Ma, Nayeon Kim, Jun Hyung Cha, Wonjong Kim, Chan Yeong Kim, Yong-ho Lee, Han Sang Kim, Yoon Dae Han, Dongeun Yong, Eugene Han, Sunmo Yang, Samuel Beck, Insuk Lee

**Author notes:** These authors contributed equally to this work (Junyeong Ma, Nayeon Kim). Corresponding author: Insuk Lee, Department of Biotechnology, College of Life Science and Biotechnology, Yonsei University, 50-1 Yonsei-ro, Seodaemun-gu, Seoul 03722, Republic of Korea.

## Abstract

Understanding the human gut microbiome requires comprehensive genomic catalogues, yet many lack geographic diversity and contain medium-quality metagenome-assembled genomes (MAGs) missing up to 50% of genomic regions, potentially distorting functional insights. Here, we describe an enhanced Human Reference Gut Microbiome (HRGM2) resource, a catalogue of near-complete MAGs (≥90% completeness, ≤5% contamination) and isolate genomes. HRGM2 comprises 155,211 non-redundant near-complete genomes from 4,824 prokaryotic species across 41 countries, representing a 66% increase in genome count and a 50% boost in species diversity compared to the Unified Human Gastrointestinal Genome catalogue. It enabled improved DNA-based species profiling, resolution of strain heterogeneity, and survey of human gut resistome. The exclusive use of these genomes improved metabolic capacity assessment enabling high-confidence, automated genome-scale metabolic models (GEMs) of the entire microbiota, revealing disease-associated microbial metabolic interactions. This resource will facilitate reliable functional insights into gut microbiomes.

## Introduction

Genome-resolved metagenomics transformed microbiome research by enabling genome assembly of uncultured microbes from shotgun data, producing metagenome-assembled genomes (MAGs)^1^. This advance enabled cataloging thousands of human gut microbial species, over 80% of which remain uncultured^2,3^. Despite progress in computational methods, many MAGs are incomplete or contaminated due to high community diversity in the gut microbiota, where numerous closely related species coexist, complicating accurate reconstruction^4^. Thus, quality control is critical, typically filtering MAGs as medium-quality (MQ; ≥50% completeness, ≤5% contamination) or near-complete (NC; ≥90% completeness, ≤5% contamination)^5^.

Most genomic catalogs for the human gut microbiome consist largely of MQ MAGs, with many species missing up to 50% of genomic regions. For example, the Unified Human Gastrointestinal Genome (UHGG)^3^ includes 204,938 nonredundant genomes, of which 97,157 (47.4%) are MQ MAGs. Only 3,207 of 4,644 (69%) species clusters contain NC MAGs or isolate genomes (collectively "NC genomes"). Reliance on incomplete genomes may lead to incorrect ecological or functional inferences. Missing genes in MQ MAGs can distort phylogenetic distances and pangenome analyses. Furthermore, only 56% of UHGG clusters composed solely of MQ MAGs are annotated at the species level in the Genome Taxonomy Database (GTDB)^6^, compared to 85% of clusters with at least one NC MAG. This indicates that MQ-only clusters are less likely to represent well-defined species. These limitations underscore the need for a gut microbiome catalog built entirely from high-completeness genomes to support more accurate and reliable analyses.

Another limitation of current genomic catalogs for the human gut microbiome is geographic and population bias. For example, the UHGG catalog is dominated by data from Western countries and China. The Human Reference Gut Microbiome (HRGM) catalog partially addressed this by adding thousands of MAGs from Korea, Japan, and India^2^. However, many regions remain under-represented. Incorporating MAGs from these areas may reveal new ecological and functional insights into gut microbiome.

Here, we present HRGM2, an enhanced version of the HRGM catalog, built from a greatly expanded sample set across 41 countries, including 10 previously understudied in Southeast Asia, Africa, and South America. This catalog includes only NC genomes with GUNC clade separation score (CSS) <0.45^7^. It comprises 155,211 non-redundant NC genomes from 4,824 prokaryotic species. Population-associated phyla in the human gut become more detectable with geographic expansion. Contrary to a previous report^8^, DNA-based species-level profiling using HRGM2 proved more accurate than marker-based methods, especially for samples with high species complexity. Comparisons between conspecific MQ MAGs and NC genomes demonstrated the limitations of MQ MAGs in taxonomic and functional analysis. High genome completeness also supports more reliable estimates of metabolic independence^9^ and enables high-confidence, automated genome-scale metabolic models (GEMs) for the entire gut microbiota^10^.

## Results

### Exclusive cataloging of NC genomes from the human gut microbiome in HRGM2

We collated 5,625 new shotgun metagenomic samples from 31 datasets not included in UHGG or the previous HRGM (HRGM1^2^) (**Supplementary Tables 1–2**). Contigs were generated using a *de novo* assembly pipeline and binned into MAGs through an ensemble method (**Extended Data Fig. 1a**). Combining all sources, we obtained 509,047 genomes: 316,079 from HRGM1, 3,632 isolates from the Broad Institute-OpenBiome Microbiome Library (BIO-ML) collection^11^, 115 isolates from the human Gut Microbial Biobank (hGMB)^12^, and 189,221 MAGs from the new samples. These MAGs underwent quality control (**Extended Data Fig. 1b**), including chimerism detection with GUNC^7^ and filtering of MAGs with CSS > 0.45, resulting in 413,014 genomes. We then used CheckM^13^ to assess completeness and contamination, selecting 224,351 NC genomes.

Lineage-specific workflow of CheckM occasionally underestimated completeness, particularly for uncharacterized clades, due to incorrect marker gene selection^14,15^. To address this, we re-evaluated genome quality using a universal bacterial marker set for clades with underestimated completeness^13,16^. For the phylum Patescibacteria, which lacks many universal markers, we applied the Candidate Phyla Radiation (CPR) marker set^16^. This reassessment retained 6,281 additional NC genomes, increasing the total to 230,632 (**Supplementary Table 3**). After freezing this version of the catalog and completing most analyses, CheckM2^15^ was released, which uses machine learning to address missing marker genes in some lineages. Re-binning and re-analysis using CheckM2 and MetaWRAP^17^ was infeasible due to time constraints. Instead, we used CheckM2 to validate our re-evaluation strategy. Over 90% of genomes retained through universal or CPR markers remained NC according to CheckM2 (**Extended Data Fig. 1c–d**), confirming the reliability of our reassessment.

We dereplicated 230,632 NC genomes into 4,824 species-level clusters, designating the highest-quality genome as the representative for each (**Supplementary Table 4**). After removing redundancies, we finalized HRGM2, an enhanced catalog of 155,211 non-redundant NC genomes representing 4,824 prokaryotic species (4,798 bacterial and 26 archaeal) in the gut microbiome (**Fig. 1a**). Due to difficulties in recovering rRNA regions from short-read MAGs, only 7,348 genomes (4.73%) met high-quality (HQ) criteria, which require an NC genome to contain 5S, 16S, and 23S rRNAs and tRNAs for at least 18 of 20 amino acids^5^. Specifically, only 2.08% of short-read MAGs met HQ criteria, compared to 89.6% of HiFi long-read MAGs.

**Fig. 1.**
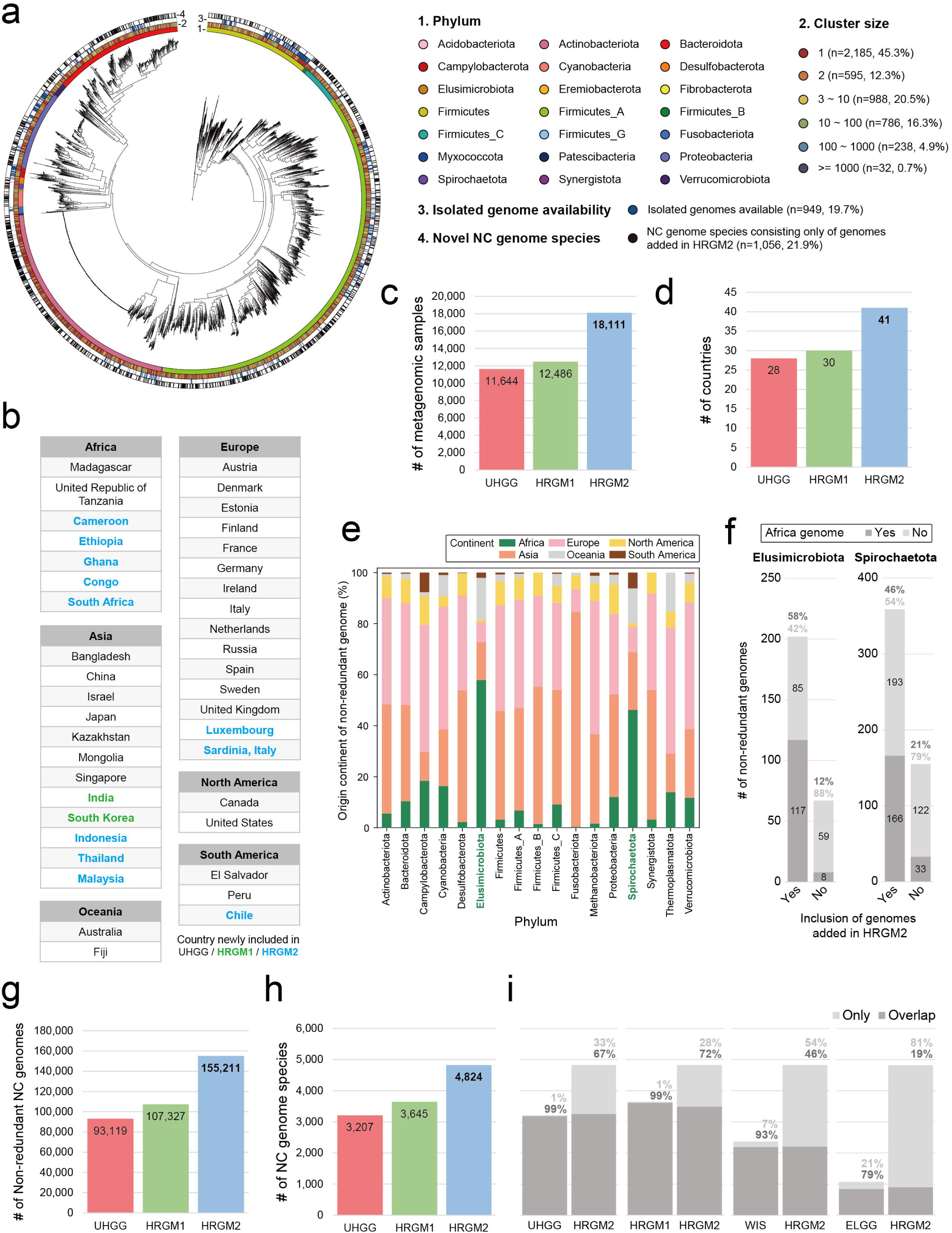
Overview of HRGM2. **a**, Maximum-likelihood phylogenetic tree representing 4,798 bacterial species in HRGM2. Color strips from inner to outer circles represent phylum, number of non-redundant genomes in the species cluster, availability of isolated genomes, and whether the species cluster consists solely of genomes newly added by HRGM2. **b**, Summary of the geographic information of the shotgun metagenomic samples used for the construction of the UHGG, HRGM1, and HRGM2. **c-d**, Comparison among UHGG, HRGM1, and HRGM2 for (c) the number of shotgun metagenomic samples from which MAGs originate and (d) the number of countries providing metagenomic samples, **e**, The proportion of continental origin of non-redundant genomes within each phylum. The plot shows only the phyla with at least 50 non-redundant genomes with known continental origin. **f**, The number and proportion of genomes originating from Africa and non-Africa according to the inclusion of added genomes in HRGM2 from the phyla Elusimicrobiota (left) and Spirochaetota (right). **g-h,** Comparison among UHGG, HRGM1, and HRGM2 for (g) the number of non-redundant near-complete (NC) genomes, and (h) the number of NC genome species. **i**, The number and percentage of NC genome species shared and not shared between HRGM2 and other human gut microbiome catalogs.

### HRGM2 improved representation of phyla enriched in samples from Africa

HRGM2 includes genomes from 18,111 shotgun metagenomic samples across 41 countries (**Fig. 1b**, **Extended Data Fig. 2**), representing 1.5 times more samples and 1.4 times more countries than UHGG and HRGM1 (**Fig. 1c–d**). This broader dataset improved representation of under-sampled regions. HRGM2 incorporated 10,855 MAGs from over 1,000 samples from Africa, a 9.3-fold increase over the 1,168 non-redundant NC genomes for Africa in UHGG and HRGM1. This expansion allowed us to observe two phyla enriched with microbial genomes from African samples: Elusimicrobiota and Spirochaetota (**Fig. 1e**). Notably, these phyla were recently found to correlate with the second principal coordinate axis of microbiota composition in African populations^18^. Abundance analysis confirmed these phyla are more prevalent and abundant in samples from Africa than in those from non-African regions, supporting genome-resolved observations (**Extended Data Fig. 3a**). The association of these phyla with samples from Africa through the inclusion of previously missing genomes in HRGM2 (**Fig. 1f**) suggests that incorporating geographically diverse samples can improve the resolution of geographic patterns in the human gut microbiome.

### HRGM2 enhances metagenomic read classification

Considering only NC genomes, HRGM2 comprises 155,211 genomes from 4,824 species, marking at least 1.45-fold and 1.32-fold increases over UHGG and HRGM1, respectively (**Fig. 1g–h**). Consequently, 99% of UHGG and HRGM1 species clusters that include at least one NC genome ("NC genome species") are also found within HRGM2 (**Fig. 1i**). Furthermore, HRGM2 captures 93% and 79% of NC genome species from other major catalogs, such as those from the Weizmann Institute of Science (WIS)^19^ and Early-Life Gut Genomes (ELGG)^20^, even though these were not incorporated into HRGM2. HRGM2 also adds 1,056 NC genome species not found in UHGG. In contrast, only 3,207 of 4,644 UHGG species (69%) contain NC genomes. Notably, among 1,437 species in UHGG composed only of MQ MAGs, 66% include a single MAG, which may not reliably support species existence in the human gut. For the 3,186 NC genome species shared by UHGG and HRGM2, HRGM2 provides, on average, 2.1 times more NC genomes per species (**Extended Data Fig. 3b**).

We evaluated taxonomic classification rates of metagenomic reads using human gut genomic catalogs. A total of 2,624 metagenomic samples not used in either catalog were aligned using Kraken2^21^ with a confidence threshold of 0.2 (**Supplementary Table 5**). Most reads mapped to both UHGG and HRGM2, but HRGM2 showed higher classification rates at all taxonomic ranks (**Extended Data Fig. 3c**). Lower rates in UHGG may result from missing regions in MQ genomes. When samples were grouped by continent, HRGM2 consistently showed higher genus-level classification rates than UHGG across all regions (**Extended Data Fig. 3d**). These results indicate that HRGM2 improves taxonomic classification rates across diverse geographic populations.

### Marker-based species-level taxonomic profiling is limited for fast-evolving species

A common use of microbiome genomic catalogs is assembly-free taxonomic profiling of metagenomic samples. Two main approaches exist: DNA-based and marker-based. DNA-based methods classify reads using full genome sequences and typically use fast k-mer aligners like Kraken2^21^, often paired with Bracken^22^ for species-level refinement. Marker-based methods classify reads using species-specific marker genes, with MetaPhlAn^8^ being a widely used tool. Custom Kraken2 databases can be easily built for any genome catalog, which is why resources like UHGG provide such databases for representative genomes^3^. In contrast, building MetaPhlAn databases requires extracting species-specific markers, making the process more labor-intensive and less commonly supported.

To compare DNA- and marker-based species-level profiling with HRGM2, we built both Kraken2 and marker gene databases. For Kraken2, we generated two versions: one using representative genomes and another using concatenated genomes per species cluster. We also constructed a species-specific marker database following the MetaPhlAn4^8^ method (**Extended Data Fig. 4a**), identifying markers for 4,209 of 4,824 species (87.3%), with over 200 markers found for 3,007 species (**Extended Data Fig. 4b**). However, the genus *Collinsella*, which shows overdispersion in HRGM2 and UHGG phylogenies, posed difficulties. Marker genes were not identified for 584 of 771 *Collinsella* species, accounting for 95% of species lacking markers (**Extended Data Fig. 4c**). This likely reflects rapid speciation, with a high density of single nucleotide variations (SNVs)^2^ but limited gene-level divergence. These results highlight a limitation of marker-based profiling for fast-evolving clades, reducing accuracy in species-level classification.

### DNA-based species profiling with HRGM2 outperforms marker-based approaches

To assess species profiling accuracy, we used simulated metagenomic samples based on HRGM2 genomes. To test the impact of species complexity, we generated samples with 40, 150, 600, 1,000, and 1,500 species, each with one or multiple strains. We evaluated two Kraken2 databases—one using representative genomes and another using concatenated genomes per species—and a MetaPhlAn4 database based on species-specific marker genes. Since longer genomes increase the chance of read alignment^23^, we examined the effect of genome size normalization on profiling accuracy. We also tested the impact of excluding reads with a Kraken confidence score below 0.2, given that ambiguous reads can skew abundance estimates^24^.

In terms of computational efficiency, Kraken2-based pipelines were about twice as fast as MetaPhlAn4 but required more memory (**Fig. 2a**). For species profiling accuracy, MetaPhlAn4 showed the highest precision, while Kraken2 pipelines using representative genomes or no confidence threshold had the lowest. However, the Kraken2 pipeline with concatenated genomes and a confidence threshold reached precision similar to MetaPhlAn4, especially in samples with over 600 species (**Fig. 2b**, **top**). These results suggest that incorporating species pangenomes and reducing false positives are key to improving precision in Kraken2-based species classification, particularly under high species complexity.

**Fig. 2.**
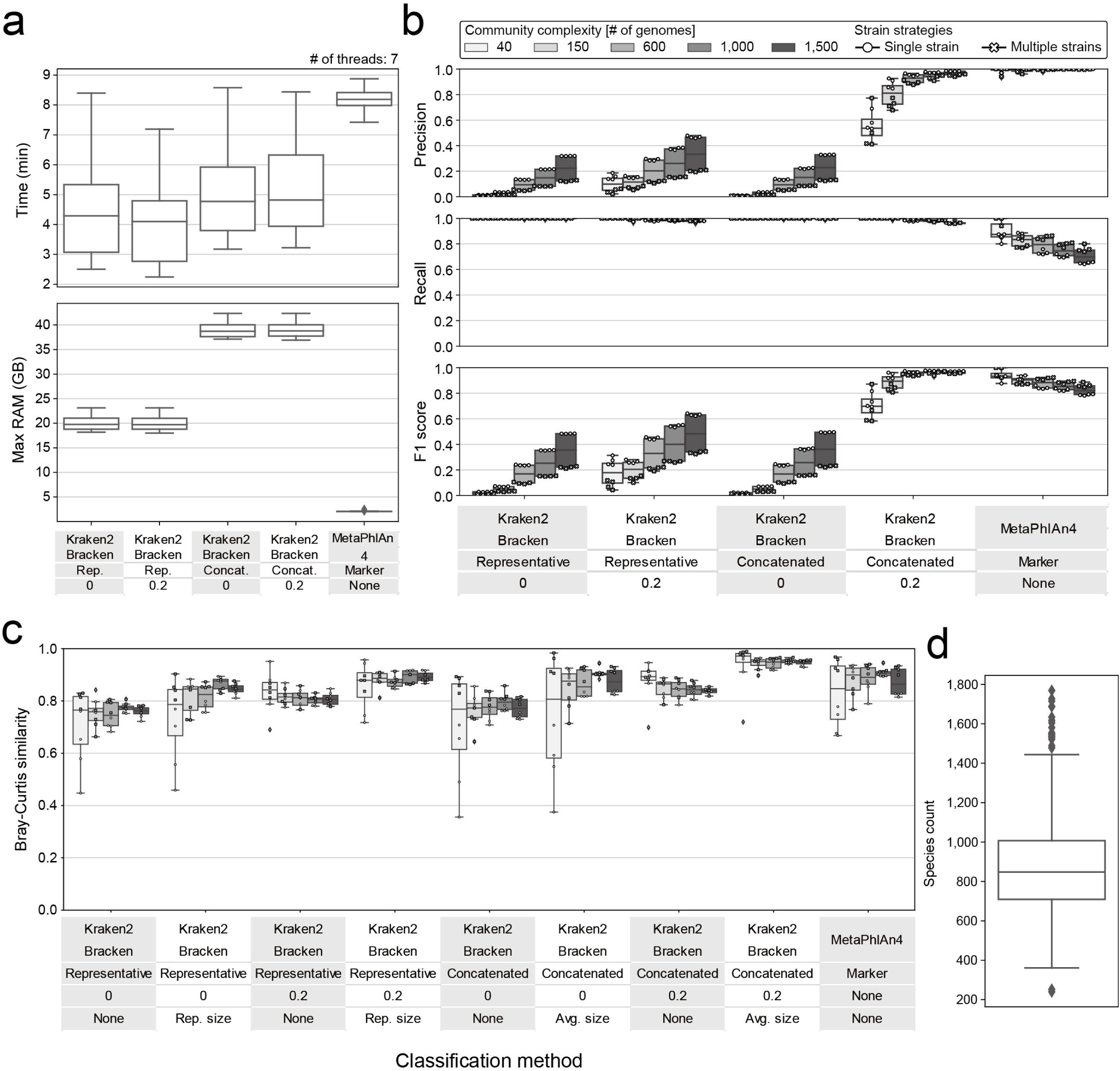
Benchmarking of taxonomic classification methods based on HRGM2 for human gut microbiota. **a-c**, Comparative analysis of taxonomic classification methods from three perspectives: (a) computational resources, (b) species detection, and (c) species abundance. Classification methods are represented by three layers in (a) and (b), and by four layers in (c). From the topmost layer, each layer represents the tool used for taxonomic classification, the database version, the confidence score, and the method of species size normalization, respectively. In (b) and (c), the color of the box indicates the number of genomes used to create the simulation data (community complexity), and the shape of the dot represents how many genomes a species is represented by (strain strategies). **d**, Number of HRGM2 species in 1,160 metagenomic samples from healthy human guts. The best pipeline according to our analysis was used, and to consider false positives, we assumed that a species was present in the sample if its normalized relative abundance was greater than 1e-06. In Fig. 2, box lengths represent the interquartile range of the data, and whiskers extend to the lowest and highest values within 1.5 times the interquartile range from the first and third quartiles, respectively. The center bar represents the median. All the outliers are shown in the plots.

Unlike precision, all Kraken2-based pipelines achieved near-perfect recall, while MetaPhlAn4 showed much lower recall, which declined with increasing species complexity (**Fig. 2b**, **middle**). This indicates that using full genome sequences provides clear advantages over marker genes for species retrieval. The reduced recall in MetaPhlAn4, especially in complex communities, may result from more species lacking specific marker genes, such as those in the genus *Collinsella*.

The F1 score, balancing precision and recall, showed that the Kraken2 pipeline with concatenated genomes and a confidence threshold performed best in samples with over 600 species, while MetaPhlAn4 performed better in simpler communities with 40 or 150 species (**Fig. 2b**, **bottom**). Strain complexity had a minimal impact. Bray-Curtis similarity further supported the superior performance of the Kraken2 pipeline with concatenated genomes, genome-size normalization, and a confidence threshold in HRGM2-based profiling (**Fig. 2c**). Analysis of 1,160 healthy gut samples (**Supplementary Table 2**) showed that 89% contained more than 600 species (**Fig. 2d**). Given this high complexity, the Kraken2 pipeline with these enhancements provides the most accurate species profiles. These findings underscore the need to consider taxonomic complexity when evaluating species profiling methods.

### HRGM2 enhanced resolution of strain heterogeneity in the gut microbiota

Global-scale microbiome studies have highlighted the value of population genetics within microbial species, complementing species-level analysis^25^. While metagenotype analysis aligns metagenomic reads to reference genomes, using pre-identified SNVs enables more efficient genotyping^26,27^, allowing large-scale exploration of subspecies-level variation. SNV databases are built from genome alignments within each species. The expanded set of NC genomes in HRGM2 increases detectable intraspecies SNVs, improving resolution of strain-level heterogeneity^28^. As most tools recommend NC over MQ genomes for better SNV detection^37,38,28^, we compared species counts eligible for SNV-based analysis at varying NC genome thresholds. HRGM2 consistently retained more qualifying species than UHGG (**Extended Data Fig. 5a**). Among species with ≥10 non-redundant NC genomes, HRGM2 detected more SNVs in 82% of cases (**Extended Data Fig. 5b**), demonstrating improved resolution of subspecies-level variation.

To examine subspecies genetic structure across geographic regions, we analyzed 1,787 publicly available gut metagenome samples (1,169 from Asia—India, Japan, China; 618 from Europe/US—Austria, Italy, Germany, France, USA; **Supplementary Table 6**). Using GT-Pro^27^, we performed SNV genotyping for 412 species with ≥50 NC genomes and sufficient prevalence, then calculated pairwise phylogenetic distances between strains from Europe/US and Asia. Geographic stratification was evaluated using the PERMANOVA test, with significance defined as *p-value* < 0.01 and pseudo-F > 30. This analysis identified 83 species (∼20%), mostly from Firmicutes_A, with clear subspecies-level geographic separation (**Extended Data Fig. 5c**, **Supplementary Table 7**). Phylogenetic trees based on SNVs for the top 20 stratified species are shown in **Extended Data Fig. 5d**. These results show that HRGM2 facilitates more comprehensive exploration of subspecies genetic structure linked to host-associated factors.

### HRGM2 enhanced functional annotations of microbial proteins

To construct a protein catalog for the human gut microbiome, we identified 549,278,140 coding sequences from 230,632 NC genomes. Redundancy was reduced via clustering at 100%, 95%, 90%, 70%, and 50% identity thresholds, resulting in ∼87.0, 18.2, 12.4, 6.5, and 3.9 million clusters, respectively.

Using HRGM2 genomes and proteins, we performed functional annotation (**Extended Data Fig. 6a**). We applied eggNOG-mapper^29^, an orthology-based approach, for KEGG Orthology (KO)^30^, Gene Ontology (GO)^31^, Protein families (PFAM)^32^, Carbohydrate-active enzyme (CAZyme)^33^ and Cluster of Ortholog Groups (COG)^34^ annotations. Due to low GO coverage, we used deep learning-based DeepGOPlus^35^, which increased GO annotation by 5.2-, 18.7-, and 5.9-fold for biological process, cellular component, and molecular function annotation, respectively, compared to eggNOG-mapper (**Fig. 3a**). Assuming species within the same taxonomic clade share similar functions, we evaluated annotation accuracy by measuring taxonomic homogeneity of genomes clustered by GO term profiles. GO profiles from DeepGOPlus showed higher accuracy than those from eggNOG-mapper (**Fig. 3b**).

**Fig. 3.**
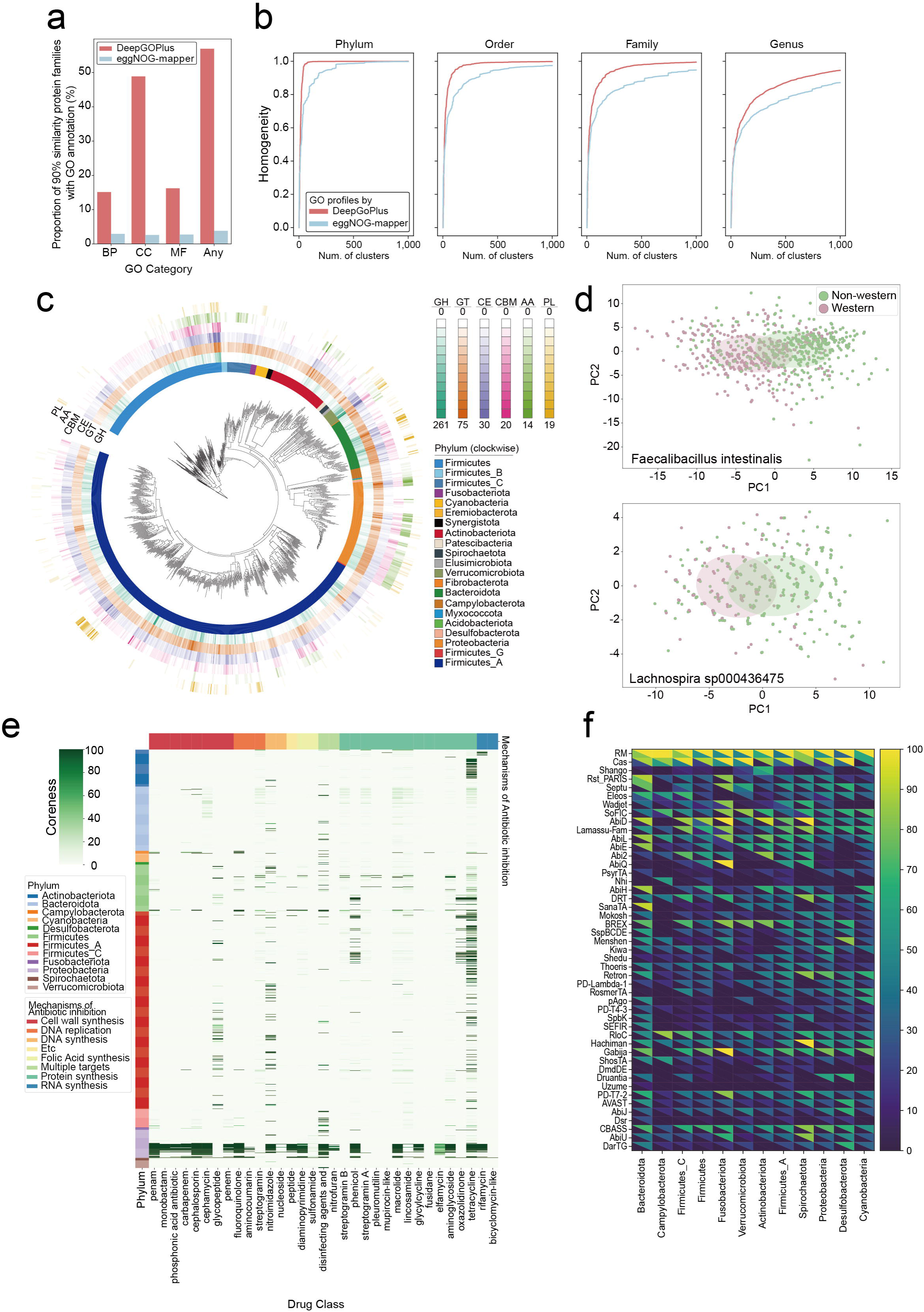
Overview of the functional landscape of the human gut microbiome. **a**-**b**, Comparison of Gene Ontology (GO) annotations by eggNOG-mapper and DeepGOPlus. (a) Annotation rates for each GO category (BP: Biological Process, CC: Cellular Component, MF: Molecular Function, Any: Any GO terms). (b) Taxonomic homogeneity of clusters resulting from agglomerative clustering based on GO annotations from either DeepGOPlus or eggNOG-mapper across different taxonomic levels. **c**, Phylogenetic tree of genera in HRGM2, indicating the average number of CAZyme families in genomes belonging to each genus. CAZyme classes; GH: Glycoside Hydrolase, GT: Glycosyl Transferase, CE: Carbohydrate Esterase, CBM: Carbohydrate Binding Module, AA: Auxiliary Activity, PL: Polysaccharide Lyase. **d**, PCoA of two species with the most distinct CAZyme profiles between Western and non-Western countries. **e**, Overview of antibiotic resistance gene in human gut microbiome. Coreness, which refers to the percentage of conspecific genomes within a species that have a resistance to each antibiotics. **f**, Overview of a defense system for viruses. Percentage of presence of each defense system as core or accessory genomes per phylum. At the bottom of each cell in the heatmap is the percentage of species where the defense system is present in their core genome, while at the top is the percentage of species where the defense system is present in their accessory genome.

CAZyme play a crucial role in the gut by breaking down various carbohydrates, including dietary fiber, starches, and polysaccharides, into various short-chain fatty acids (SCFAs) that impact host health^36^. Additionally, some CAZymes can degrade mucin, which protects the intestinal epithelium from pathogens. CAZyme annotations, performed using dbCAN3^37^, revealed that Verrucomicrobiota and Bacteroidota had the most CAZyme families on average (**Extended Data Fig. 6b**). Distribution varied more at lower taxonomic levels (**Fig. 3c**). The genus with the most glycoside hydrolases (GHs), enzymes critical for carbohydrate degradation^38^, was identified (**Extended Data Fig. 6c**); in Firmicutes, genera from the Paenibacillaes order exhibited high GH abundance, while in Firmicutes_A, *Robinsoniella*, *Murimonas*, and *Hungatella* were dominant.

Given dietary influence on CAZyme activity^39^, we hypothesized diet-associated divergence within species. Indeed, we identified several species with distinct CAZyme profiles between Western and non-Western countries (**Fig. 3d**, **Extended Data Fig. 6d**). *Faecalibacillus intestinalis* showed the strongest divergence, with non-Western strains enriched in GH1, GH140, GH128, and GH55—enzymes linked to fiber degradation^40^ (**Extended Data Fig. 6e**).

### HRGM2 enables reliable surveys of human gut resistome and phage defense systems

Gut bacteria have evolved defense systems against antibiotics and viruses, many of which spread via horizontal gene transfer (HGT)^40,41^. HRGM2, composed of only NC genomes, enables reliable analysis of resistance gene distribution across core and accessory genomes. Genes in core genomes suggest vertical inheritance, while those in accessory genomes indicate HGT.

Using RGI^41^, we predicted resistance genes in 1,042 species with ≥10 NC genomes, focusing on phyla with ≥5 species. We quantified the proportion of genomes per species carrying specific resistance genes (**Fig. 3e**, **Supplementary Table 8**). Results confirmed previous findings^42,43^: Proteobacteria species often carry multiple resistance genes, especially for antibiotic efflux.

Actinobacteriota phylum showed lineage-specific resistance to tetracycline and rifamycin, with *Bifidobacterium* (e.g., *B. adolescentis*) carrying rpoB mutations for rifamycin resistance^44^, and *Collinsella* harboring Tet(Q) for tetracycline resistance. In Bacteroidota phylum, most resistance genes were found in accessory genomes.

We identified species with multiple resistance types in core genomes, suggesting high transmission potential. These were mostly from *Escherichia*, *Salmonella*, *Klebsiella*, and *Enterobacter*, all of which belong to Proteobacteria phylum. Outside this phylum, *Staphylococcus epidermidis* (MRSE)^45^ and *S. aureus* (MRSA)^46^ from Firmicutes phylum carried 10 and 6 resistance types, respectively, while BX12 sp014333425 from Firmicutes_A carried five.

NC genome-based HRGM2 also enabled reliable profiling of antibiotics with the highest resistance gene prevalence. Tetracycline resistance was most common, detected in 561 (53.8%) out of 1,042 species carrying resistance genes, followed by lincosamide and macrolide resistance in 278 (26.7%) species.

Lastly, we examined phage defense systems. Most, excluding RM and Cas systems, were located in accessory genomes (**Fig. 3f**). Exceptions included the Shango system in Actinobacteriota and the Rst_PARIS system in Bacteroidota core genomes. Other systems were broadly distributed across phyla.

### MQ genomes may result in unreliable metabolic profiles of the gut microbiota

A key distinction of HRGM2 over UHGG is its exclusive inclusion of NC genomes. In UHGG, only 66% of 4,644 species include NC genomes, and 55% of the 204,938 non-redundant genomes are MQ (**Extended Data Fig. 7a**). By median, 55% of genomes per species cluster are MQ (**Extended Data Fig. 7b**), indicating lower quality in representative and pangenomes compared to HRGM2. Incomplete MQ genomes may cause false negatives in functional profiling. To evaluate this, we analyzed 327 species represented only by MQ genomes in UHGG but with at least one NC genome in HRGM2. Aligning conspecific representative genomes showed that 89% of MQ genomes overlapped with NC genomes, but 26% of NC genomes were absent from conspecific MQ genomes (**Fig. 4a**). Clustering predicted proteins from each representative genome yielded similar results (**Fig. 4b**). Both genome and protein analyses revealed missing content in MQ genomes relative to NC counterparts.

**Fig. 4.**
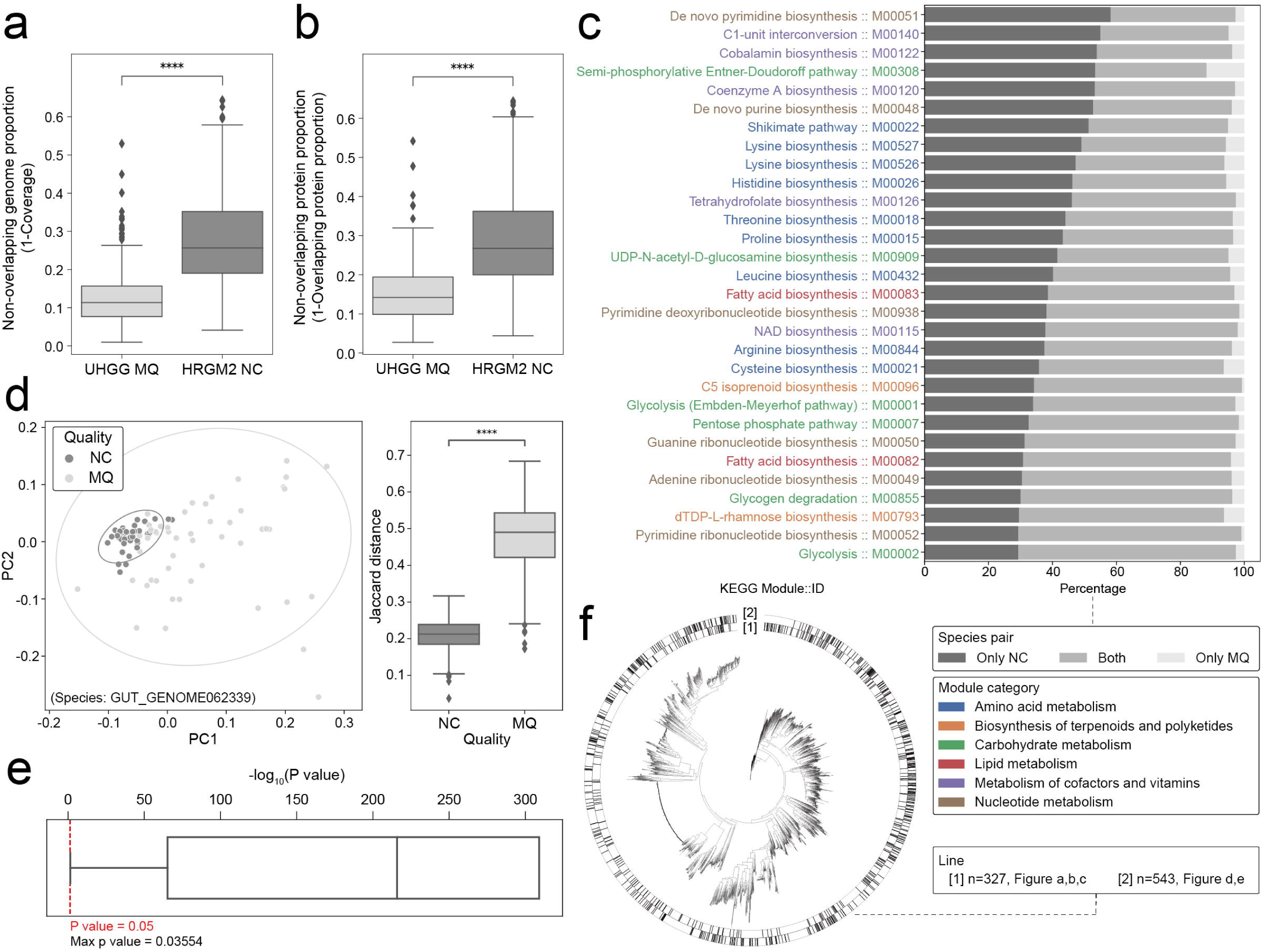
The limitations of MQ genomes in functional profiling. **a**-**b**, Comparison of non-overlapping (a) genome and (b) protein proportion for conspecific MQ and NC genomes. Differences were evaluated with two-sided Wilcoxon signed-rank test (P = (a) 5.35e-42; (b) 1.06e-39). **c**, Top 30 KEGG modules with a high percentage detected only in NC genomes. **d**, PCoA plot of the functional composition profiles of MQ and NC genomes (left) and the functional distance between genomes within the same quality group (right) for the species ‘GUT_GENOME062339’ in UHGG. Ellipses in PCoA represent a 95% confidence interval. A one-tailed Mann-Whitney U test was performed to confirm if the functional distances between NC genomes were significantly smaller than between MQ genomes (P < 1e-300). **e**, *P*-values of the 543 species evaluating the functional distance difference as in (**d**, right), calculated using the same one-tailed Mann-Whitney U test. To take the logarithm of *P*-values, *P*-values of 0 were replaced with the smallest non-zero *P*-value (9.11e-310). The red dotted line represents a *P*-value of 0.05. **f**, Position of the UHGG species used in the analyses on the UHGG phylogenetic tree. The inner and the outer black color strips represent the positions of the 327 species used in (**a-c**) and the 543 species used in (**d**, **e**), respectively. For boxplots in Fig. 4, box lengths represent the interquartile range of the data, and whiskers extend to the lowest and highest values within 1.5 times the interquartile range from the first and third quartiles, respectively. The center bar represents the median. All the outliers are shown in the plots. ****, *P* < 1e-04.

To assess how missing regions in MQ genomes affect functional analysis, we predicted KEGG modules^47^ for each representative genome and identified functions found exclusively in NC genomes of conspecific pairs. Core biosynthesis modules for amino acids, cofactors, vitamins, and nucleotide metabolism were often absent in MQ genomes (**Fig. 4c**), suggesting false negatives in metabolic profiling.

We then compared functional profiles of MQ and NC genomes within species, analyzing 543 UHGG species with ≥10 non-redundant genomes of each type. Functional distance, based on gene clusters, was significantly higher among MQ genomes, which showed more variability, while NC genomes had consistent profiles (**Fig. 4d**). This pattern was observed across all tested species (**Fig. 4e**) and aligns with prior findings from the GTDB database^48^. These results indicate that functional variation among NC genomes reflects biological differences between strains, whereas variation among MQ genomes is mainly technical. The phylogenetic distribution of analyzed species shows that this bias is not limited to specific clades (**Fig. 4f**). These findings support the need for exclusive use of NC genomes in gut microbiome reference catalogs.

### HRGM2 provides a reliable landscape of gut microbiota metabolic capacities

Most prokaryotic species in the human gut remain uncultured, partly due to metabolic dependence. Understanding metabolic capacities helps clarify microbial interactions and colonization. A recent study estimated "metabolic independence" using 33 KEGG modules, mostly related to essential metabolite biosynthesis, and enriched in species that successfully colonize after fecal microbiota transplantation^9^. Species with most of these modules were classified as high metabolic independence (HMI), while those lacking them were termed low metabolic independence (LMI). Using this framework, we assessed gut microbial species in HRGM2.

As shown earlier, MQ genomes often underestimate metabolic capacity, frequently misclassifying species as LMI. In contrast, HRGM2, composed solely of NC genomes, reduces such underestimation in profiling with the 33 KEGG modules. Among 4,824 HRGM2 species, 751 (15.6%) were HMI and 388 (8.0%) LMI. Compared to HRGM2, UHGG species, many of which are based on MQ genomes, had fewer KEGG modules, resulting in fewer HMI and more LMI classifications (**Extended Data Fig. 8a–b**). This confirms that catalogs including MQ genomes tend to underestimate metabolic capacities.

Consistent with prior studies^9^, HMI species had larger genomes than LMI species (**Extended Data Fig. 8c**). HMI species were found across more countries (**Extended Data Fig. 8d**), suggesting broader adaptation to gut environments. Metabolic independence also correlated with availability of isolated genomes—98.7% of LMI species were uncultured (**Extended Data Fig. 8e**), likely due to dependence on external metabolites.

We identified 15 HMI clades, primarily composed of HMI species such as Enterobacterales and Lachnospirales, and 7 LMI clades, five of which had placeholder names and remain poorly characterized (e.g., o RFN20, o RF39, f UBA1242, o TANB77, g UMGS1491) (**Fig. 5a**). Another LMI clade, phylum Patescibacteria, has small genomes (∼1 Mbp), is largely uncultured, and shows limited metabolic potential, suggesting a parasitic lifestyle^49^. These understudied LMI clades may also rely on parasitism, and co-culturing may aid their isolation.

**Fig. 5.**
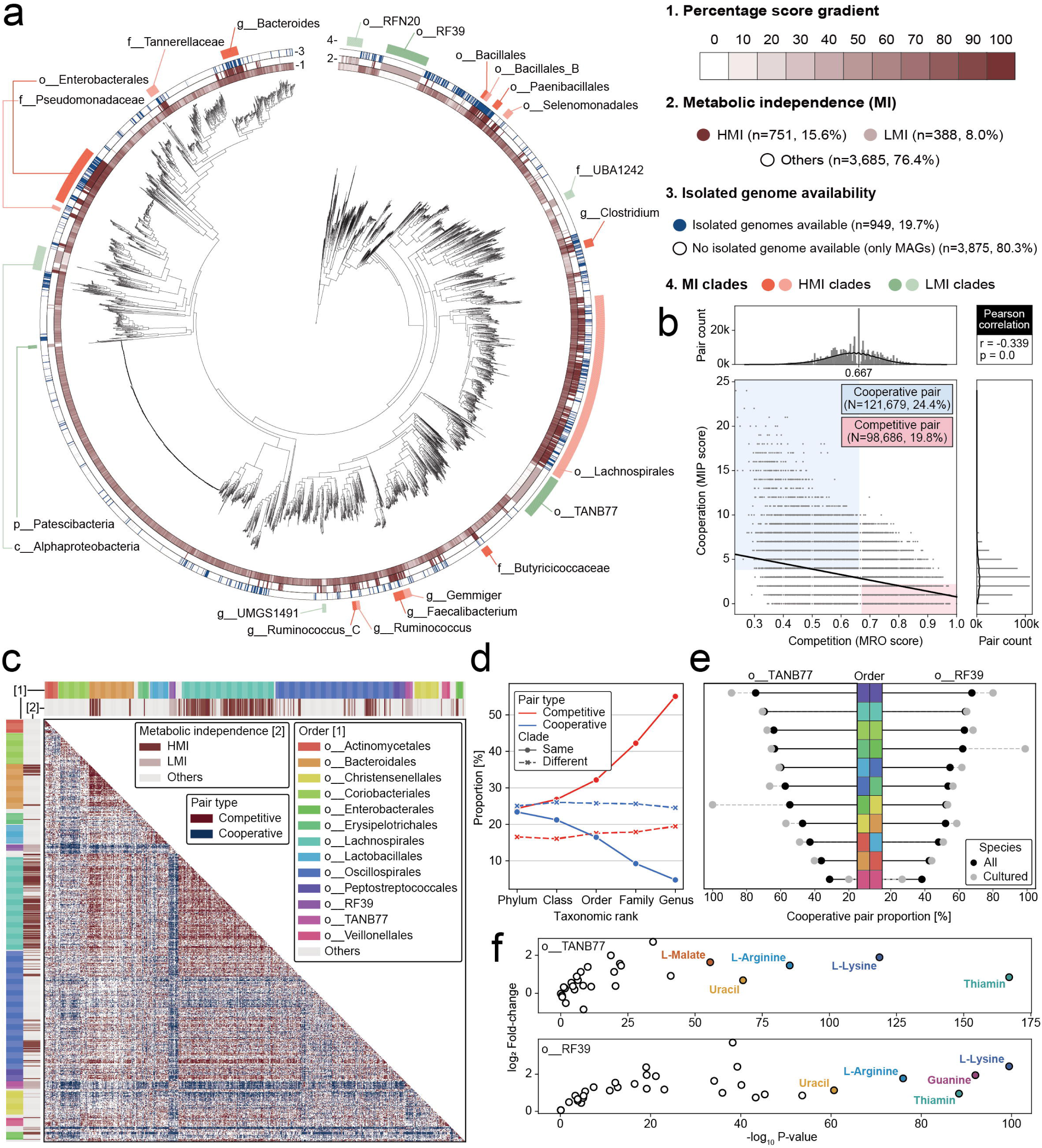
Analysis of metabolic independence and interactions in the human gut microbiota. **a**, A maximum-likelihood phylogenetic tree annotated with metabolic independence (MI) status. The inner to outer circles represent possession percentage of 33 KEGG modules used to assess metabolic independence, MI status, isolated genome availability, and designation of MI clades. **b**, A scatter plot showing distribution of metabolic resource overlap (MRO) and metabolic interaction potential (MIP) scores across all species pairs (n=499,500) among the top 1,000 species prevalent in healthy human guts globally. The blue and pink regions indicate cooperative (n=121,679, 24.4%) and competitive (n=98,686, 19.8%) pairs, respectively. A linear regression line highlights the relationship between MRO and MIP scores, with correlation coefficient and two-sided *P*-value determined by a Pearson correlation test in SciPy. **c**, A chart showing metabolic interactions, where outer color strips represent taxonomic order, and inner strips denote MI. Species are ordered by lineage, with orders containing fewer than 10 species grouped as “Others”. **d**, Proportion of cooperative and competitive pairs within the same and across different clades, with "Clade" referring to ranks from phylum to genus. **e**, Cooperative pair proportion between order TANB77 (left) / RF39 (right) and other orders is displayed. Each color represents an order, consistent with panel (**c**). "All" refers to the cooperative pair proportion across all species in an order, while "Cultured" is specific to cultured species with isolated genomes. **f**, Association between metabolite supply and the cooperative behavior of species toward TANB77 (top) and RF39 (bottom) orders. Fold-change was calculated as the ratio of the proportion of cooperative pairs that transferred a given metabolite to the LMI order to the corresponding proportion among non-cooperative pairs. *P*-values were calculated using a two-sided Fisher’s exact test. Each dot represents a metabolite, and for each LMI order, the five most significant metabolites based on *p*-values are highlighted in distinct colors.

### HRGM2 supports genome-scale metabolic modeling for entire gut microbiota

Understanding metabolic interactions among microbes may enable modulation of microbiome composition and function, influencing host physiology^10^. Modeling these interactions is becoming essential for personalized nutrition and healthcare. Metabolic modeling primarily relies on GEMs, which represent an organism’s full metabolic network^50^. GEMs were initially reconstructed for culturable species with complete genomes but have recently been developed for thousands of gut bacterial strains^51,52^. The rise of nearly complete MAGs has enabled automated GEM reconstruction^53^, though high genome completeness remains critical. While automated tools can process medium- or low-quality MAGs, low completeness reduces model quality and limits downstream analysis. Studies show that GEMs from NC MAGs contain, on average, 24.6% more genes, 12.8% more reactions, and 10.3% more metabolites than those from MQ MAGs^54^. Moreover, lower completeness increases the proportion of gap-filled reactions^55^, highlighting the importance of genome quality for reliable GEMs.

By using only NC genomes, HRGM2 serves as an ideal resource for MAG-based genome-scale metabolic modeling, enabling reliable GEM construction through automated pipelines. Using CarveMe^56^, we generated GEMs for 4,824 representative species genomes. To assess the effect of genome completeness, we also built GEMs for 327 conspecific MQ-NC genome pairs from UHGG and HRGM2 and compared their metabolic content. While most GEM elements (metabolites, reactions, gene-associated reactions) in MQ models were also present in NC models, NC GEMs consistently included more unique elements not captured by MQ models (**Extended Data Fig. 8f**). Notably, gene-associated reactions showed the largest difference in the proportion of unique elements. These results show that MQ-derived GEMs are lower in quality and may cause false negatives in downstream functional analyses.

Using GEMs from representative genomes, we analyzed metabolic interactions among gut microbes with SMETANA^57^ in "global" mode. This tool estimates metabolic resource overlap (MRO), indicating competition potential based on nutrient requirements, and metabolic interaction potential (MIP), indicating cooperation potential based on exchangeable metabolites. To validate predictions, we used 31 gut bacterial strains tested for their ability to decolonize *Klebsiella pneumoniae* Kp-2H7 through carbon competition^58^. Among them, 18 strains (F18-mix) showed stronger decolonization, while 13 strains (F13-mix) showed weaker effects. We expected F18-mix strains to display lower cooperation potential and higher competition potential with Kp-2H7 than F13-mix strains. Although not statistically significant due to small sample size, the trend matched expectations: F18-mix strains exhibited lower MIP and higher MRO with Kp-2H7 (**Extended Data Fig. 8g**). These results support the utility of HRGM2-based GEMs for predicting microbial metabolic interactions.

We analyzed metabolic interactions among 1,000 species most prevalent in healthy human guts across various countries (**Supplementary Table 9**). A significant negative correlation was found between MRO and MIP (**Fig. 5b**). Based on MRO and MIP values, 121,679 species pairs with MIP > 3 and MRO < 0.667 were defined as cooperative (24.4%), while 98,686 pairs with MIP < 3 and MRO > 0.667 were defined as competitive (19.8%).

We then mapped cooperative and competitive interactions among the 1,000 species **(Fig. 5c)**. HMI species tended to compete with other HMI species, and LMI species with other LMI species. Species from the same order were predominantly competitive with each other. Analysis of pairwise interactions within and across clades (**Fig. 5d**) showed that pairs within the same clade were mostly competitive, consistent with earlier studies^57^. This pattern intensified at finer taxonomic levels, reflecting increased metabolic similarity among closely related species.

The two LMI clades, RF39 and TANB77 orders, were generally cooperative with other clades—likely parasitic—except when paired with other LMI clades (**Fig. 5c)**. Cooperative pair proportions were highest with Peptostreptococcales, Lachnospirales, and Coriobacteriales, and lowest with Veillonellales (**Fig. 5e**). These patterns remained similar when restricted to cultured species, except for Veillonellales, suggesting that clades with high cooperation may aid in culturing RF39 and TANB77. Notably, Peptostreptococcales and Coriobacteriales, despite high cooperation rates, had very few HMI species (2/29 and 0/73, respectively) and contained only 58% and 61% of the 33 KEGG modules on average. In contrast, Bacteroidales, which includes the HMI-rich *Bacteroides* genus, showed low cooperation.

We hypothesized that specific metabolites facilitate cooperation with RF39 and TANB77. Using SMETANA’s "detailed" mode, we identified thiamin, L-lysine, L-arginine, and uracil as commonly supplied metabolites (**Fig. 5f**). Thiamin acts as a coenzyme in energy metabolism^59^; L-lysine supports energy metabolism^60^ and cell wall synthesis^61^; L-arginine provides carbon and nitrogen for metabolism and biofilm regulation^62^; uracil, essential for nucleic acid metabolism, may be scavenged via the salvage pathway^63^. These results suggest that RF39 and TANB77 depend on these metabolites but lack the biosynthetic capacity to produce them.

### HRGM2 GEMs reveal disease-associated microbial metabolic interactions

To demonstrate the utility of HRGM2 GEMs for studying gut microbial functions, we explored the metabolic interaction patterns within disease-associated microbial communities. We focused on two diseases—Crohn’s disease (CD) and colorectal cancer (CRC)—with bacterial species identified through an optimized machine learning pipeline^64^. We selected seven CD-associated species (*Klebsiella pneumoniae, Allisonella pneumosintes, Lactiplantibacillus plantarum, HGM08974 sp900766555, Dialister invisus, Enterococcus faecalis,* and *Pseudescherichia sp002298805*) and 11 CRC-associated species (*Fusobacterium animalis, Prevotella intermedia, Fusobacterium sp000235465, Allisonella pneumosintes, Gemella morbillorum, Fusobacterium_C necrophorum, Fusobacterium polymorphum, Porphyromonas_A somerae, Peptostreptococcus stomatis, Porphyromonas sp900539765,* and *Parvimonas micra*). Using SMETANA, we mapped metabolic interactions at both the community and pairwise levels. Interestingly, "cooperative interactions" were enriched in the CD-associated microbial community, whereas "competitive interactions" dominated in the CRC-associated community (**Fig. 6a**). These patterns were consistently observed at the pairwise level (**Fig. 6b**).

**Fig. 6.**
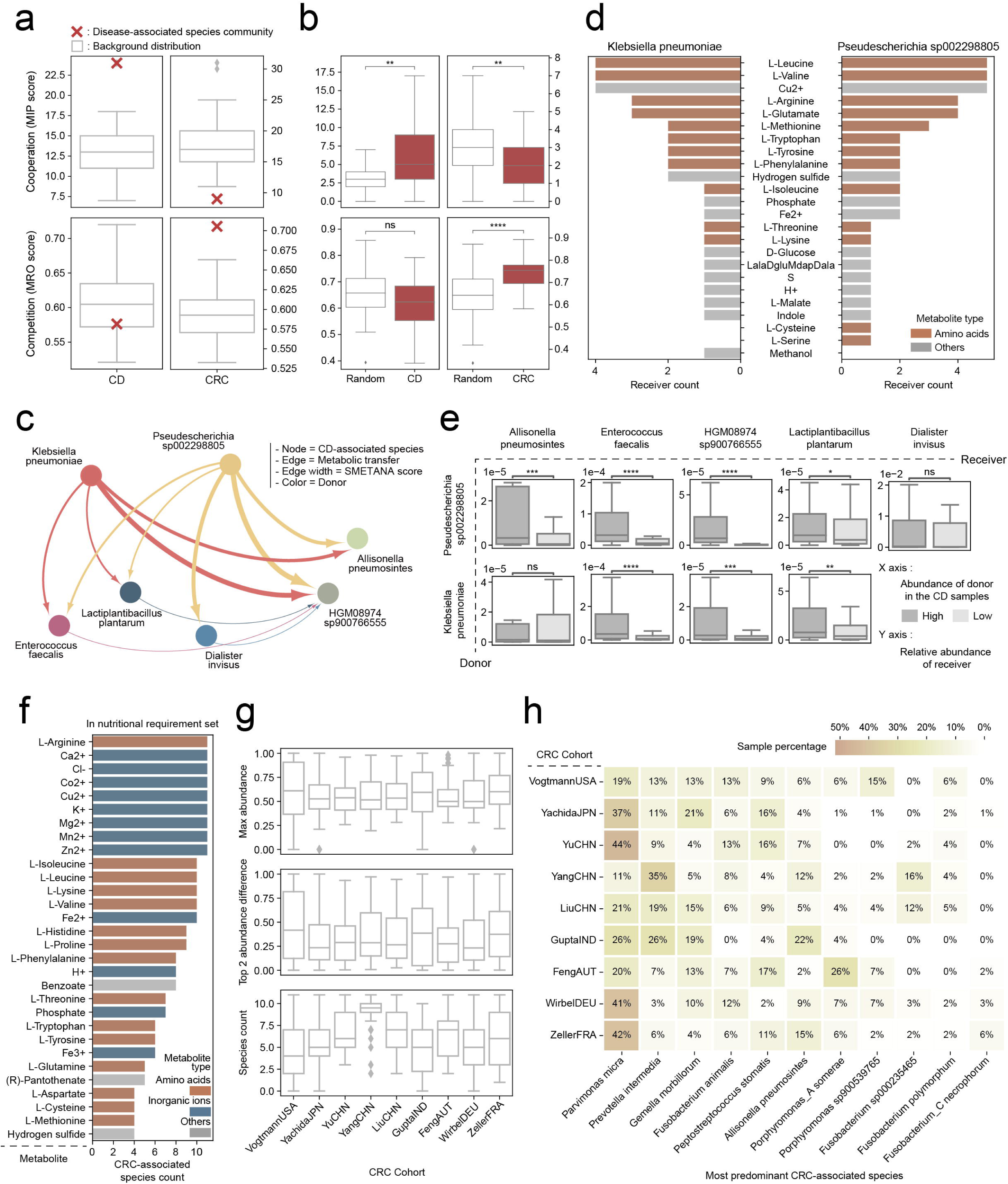
Distinct metabolic interaction patterns among CD- and CRC-associated species. **a-b**, MIP and MRO distributions of CD- and CRC-associated species at the (a) community-wise and (b) pairwise levels (n = 21 for CD and n = 55 for CRC), white boxplots show distributions from random communities (n = 100) or pairs (n = 42 for CD and n = 110 for CRC) for comparison. **c**, Metabolic interaction pattern among CD-associated species: node, edge, edge width, and color represent species, metabolic transfer, SMETANA score, and donor identity, respectively. **d**, Key metabolites delivered to the receiver species by the two main donors, *Klebsiella pneumoniae* (left) and *Pseudescherichia sp002298805* (right). **e**, Receiver species abundance according to donor abundance (high, n = 112 or low, n = 113) in CD samples. **f**, Top 30 metabolites commonly required by CRC-associated species. **g**, Distribution of the relative abundance of the top dominant species (top), abundance gap between the first and second dominant species (middle), and the number of detected CRC-associated species per sample (bottom) (VogtmannUSA, n = 52; YachidaJPN, n = 258; YuCHN, n = 45; YangCHN, n = 98; LiuCHN, n = 78; GuptaIND, n = 30; FengAUT, n = 46; WirbelDEU, n = 60; ZellerFRA, n = 53). **h**, Proportion of predominant CRC-associated species across cohort; x-axis species are ordered by prevalence across cohorts (highest to lowest). For boxplots, boxes represent the interquartile range, whiskers extend to the lowest and highest values within 1.5 times the interquartile range, and the center line indicates the median. All the outliers are shown in the plots, except (e). In (b) and (e), statistical significance was assessed using a two-sided Mann-Whitney U test. Exact P-values are as follows: (b) for CD, MIP (P = 4.431e-03) and MRO (P = 1.627e-01); for CRC, MIP (P = 1.024e-03) and MRO (P = 2.175e-10), all compared with Random; (e) for receivers (*Allisonella pneumosintes*/*Enterococcus faecalis*/*HGM08974 sp900766555*/*Lactiplantibacillus plantarum*/*Dialister invisus*): donor *Pseudescherichia sp002298805* = 4.977e-04/2.760e-08/4.021e-28/4.539e-02/1.065e-01; donor *Klebsiella pneumoniae* = 4.446e-01/6.690e-08/2.177e-04/5.750e-03. ns: not significant, P > 0.05; *P < 0.05; **P < 0.01; ***P < 1e-03; ****P < 1e-04.

To investigate cooperative interactions among CD-associated species, we used SMETANA in "detailed" mode. *K. pneumoniae* and *Pseudescherichia spp*. were key donors, providing metabolites to other species (**Fig. 6c**). Receivers mainly depended on donor-supplied amino acids for growth (**Fig. 6d**). The donors showed similar metabolite profiles, suggesting shared ecological roles.

We next tested whether these interactions were reflected in CD microbiota. We hypothesized that higher donor abundance would lead to increased receiver abundance due to greater metabolite availability. Among nine donor–receiver pairs, seven showed significantly higher receiver levels in high-donor samples (**Fig. 6e**). This concordance between GEM predictions and microbial abundance supports the reliability of HRGM2-based analyses. These results also suggest that disrupting *K. pneumoniae* and *Pseudescherichia spp*. may offer therapeutic strategies by interfering with metabolite exchange in CD-associated communities.

In contrast, CRC-associated species showed a competitive metabolic profile. To identify metabolites driving this competition, we extracted species-specific growth requirements from SMETANA’s MRO calculation. Shared dependencies indicated competitive bottlenecks, with species competing mainly for amino acids and inorganic ions (**Fig. 6f**). Notably, *F. animalis* and *F. polymorphum*, subspecies of *F. nucleatum*, are known to ferment amino acids into butyrate and acetate^65^.

We hypothesized that sustained competition would favor dominance by the most competitive species. In CRC microbiota, the top species showed a median relative abundance of 0.53, with a 0.27 median gap between the first and second most abundant species, and the median count of CRC-associated species detected per sample was six (**Fig. 6g**). This indicates that one species often dominates over half the CRC-associated community. As predicted, species dominance was consistent across samples.

Further analysis showed variation in the dominant species across cohorts, with *P. micra* most frequently dominant (**Fig. 6h**), suggesting a strong competitive advantage over other CRC-associated species.

## Discussion

A complete reference genome set for commensal microbes is key to advancing human microbiome research. Despite years of effort to build a comprehensive gut microbiome catalog, challenges such as geographic bias and limited genome quality persist. In this study, we addressed these issues by collating metagenomic data from diverse geographic regions and including only NC genomes to construct HRGM2.

We expanded sample coverage to 41 countries—a 1.5-fold increase over UHGG, which includes 28 countries but is biased toward Western regions and China, limiting global microbial representation. Our broader dataset improved detection of two phyla associated with samples from Africa, Elusimicrobiota and Spirochaetota, previously underrepresented due to limited sampling in Africa. These observations suggest that sampling bias can obscure taxonomic diversity and that targeting underrepresented regions is key to further improving HRGM2.

Another key improvement in HRGM2 is the exclusive use of NC genomes, which strengthens functional analysis of the microbiome. Existing catalogs often include MQ genomes, whose missing regions can cause false negatives and misrepresent functional profiles. We presented several case studies showing the benefits of using only NC genomes, including: (1) more complete surveys of resistome and phage defense systems, (2) more reliable genome-level functional profiles, (3) reduced false identification of LMI microbes, and (4) robust metabolic modeling through automated GEM reconstruction. These results support that catalogs based solely on NC genomes provide more accurate functional insights into the gut microbiome.

To support HRGM2-based taxonomic profiling, we developed custom databases for both DNA-based and marker-based methods. Contrary to a previous report^8^, DNA-based profiling using HRGM2 outperformed marker-based methods in samples with high species complexity (>600 species). In over a thousand healthy individuals, 89% harbored more than 600 species, supporting Kraken2/Bracken as the preferred pipeline for HRGM2-based species profiling. Several factors may explain this discrepancy. Marker genes are difficult to define for fast-evolving species like *Collinsella*, resulting in unresolved taxa. The previous study used a broad genome database for Kraken2, while we used a gut-specific one, reducing false positives. We also applied a strict confidence threshold (0.2), improving accuracy. Lastly, the earlier report did not evaluate profiling at high species complexity, which our analysis shows is common in the human gut. Therefore, although both approaches are available, we recommend Kraken2/Bracken with genome size normalization and a 0.2 confidence cutoff for accurate species profiling with HRGM2.

HRGM2 has several limitations. First, although all genomes meet NC criteria, many do not meet HQ standards. While HRGM2 is higher quality than existing catalogs, a truly complete reference will require MAGs generated by long-read, high-accuracy sequencing platforms. Second, some geographic populations remain underrepresented. We added MAGs from Africa, Southeast Asia, and South America, but regions like West Asia, North Africa, and parts of Central and South America are still under-sampled, requiring further efforts. Finally, HRGM2 includes only prokaryotic genomes. As interest grows in gut-associated viruses and fungi, a more comprehensive catalog will need to incorporate genome sequences from these organisms.

Given the continued growth of public metagenomic datasets and advances in MAG reconstruction methods, we plan to update HRGM every 2–3 years.

## Acknowledgements

This research was supported by the National Research Foundation funded by the Ministry of Science and ICT (2022M3A9F3016364, 2022R1A2C1092062 to I.L.). This work was also supported by the Technology Innovation Program (20022947) funded by the Ministry of Trade Industry & Energy (MOTIE, Korea). The work was supported in part by Brain Korea 21(BK21) FOUR program.

## Author Contributions Statement

JM, NK, and IL conceived this study. JM and NK constructed the catalog and performed bioinformatics analysis. JHC, WK, and SB contributed to bioinformatic analysis. CYK provides technical and scientific advice. Y.L., H.S.K., Y.D.H., D.Y., and E.H. contributed sequencing data generated from unpublished studies. SY constructed the web server. IL supervised the project. JM, NK, and IL wrote the manuscript. All authors read and approved the final manuscript.

## Online Methods

### Sample collation and metagenome assembly

We collated 5,625 shotgun metagenomic samples from 31 human gut metagenomic datasets, which were not included in the Unified Human Gastrointestinal Genome (UHGG)^3^ and the previous version of Human Reference Gut Microbiome (HRGM1)^2^. For public data collation, we queried the European Nucleotide Archive (ENA) using keyword combinations including: (1) country name, (2) ‘gut’, ‘fecal’, or ‘stool’, (3) ‘metagenome’ or ‘whole’, and (4) ‘human’. Keywords were joined with ‘AND’ operators for the search. To address gaps in coverage, we manually added datasets missed by this approach. We collated public datasets that had been publicly released as of February 2022. Only those with an average sequencing depth ≥3 Gbp were retained; for countries unrepresented in prior catalogs, datasets ≥1 Gbp were included to improve geographic coverage. Samples used for the construction of UHGG and HRGM1 were excluded, as their metagenome-assembled genomes were already incorporated into HRGM2 from the previous catalogs.

Three of the 31 datasets were newly generated. Of these, 176 samples were from a Non-Alcoholic Fatty Liver Disease (NAFLD) study approved by the Institutional Review Boards (IRBs) of Severance Hospital (4-2016-0950) and Dongsan Medical Center (2017-04-040) (PRJNA1227423), 26 from a CRC study approved by the IRB of Severance Hospital (1-2022-0071) (PRJNA1226738), and 16 from a fecal banking project approved by the IRB of Severance Hospital (4-2016-0850) (PRJNA1227720). NAFLD study samples were self-collected using a commercial kit (PDXen, Seoul, Korea), frozen immediately in a portable freezer, transported to hospitals within 30 min, and stored at −80 °C. CRC stool samples were collected in cryotubes and stored at −80 °C. Fecal banking donor aliquots were suspended in PBS, cleared of debris, and stored at −80 °C with 10% glycerol. Written informed consent was obtained from all participants. Shotgun metagenomic sequencing was performed on all samples. Microbial DNA was extracted using the PowerSoil DNA Isolation Kit (Qiagen), quantified with a Qubit 3.0 fluorometer (Thermo Fisher), and 50 ng DNA was used to construct paired-end libraries. Libraries were sequenced (2 × 150 bp) on Illumina NextSeq or NovaSeq platforms to a depth of 6-30 Gb.

The adapter sequences and low-quality bases of Illumina sequencing samples were removed using Trimmomatic v0.39^66^. Next, we aligned sample reads against the human reference genome (GRCh38.p13) using Bowtie2 v2.3.5^67^ and removed the aligned reads as human contaminants. Most of the quality-controlled reads were assembled as contigs using metaSPAdes v3.14.0^68^. However, for a few samples, metaSPAdes did not terminate for unknown reasons, and metaSPAdes was only applicable to paired-end sequencing reads. For samples where metaSPAdes was unavailable, we used MEGAHIT v1.2.9^69^ to assemble reads instead **(Supplementary Table 2)**. In the case of PacBio HiFi sequencing samples, we used Minimap2 v2.18-r1015^70^ with the *-ax asm20* parameter for human contaminant removal and hifiasm-meta v0.2-r040^71^ for assembly.

We employed the ensemble approach to generate genome bins. For each sample, reads were aligned to the assembled contigs with the aligner used to remove human contaminants, and alignment results were converted to sorted BAM using SAMtools v1.10^72^. Binning was then performed using three tools: MetaBAT2 v2.15^73^, MaxBin2.0 v2.2.7^74^, and CONCOCT v1.1.0^75^. The minimum contig size for binning was set to 1,000 bp, except for MetaBAT2, which requires at least 1,500 bp, so it was set to 1,500 bp. The binning results from the three tools were combined into a more robust bin set using the bin refinement module of MetaWRAP v1.3.1^17^.

### Genome collation and bin quality control

For HRGM2, we integrated genome bins assembled from 5,625 shotgun metagenomic samples with the genomes available in the previous version of HRGM. Additionally, we integrated isolated genomes from the Broad Institute-OpenBiome Microbiome Library (BIO-ML)^11^ and a human Gut Microbial Biobank (hGMB)^12^, which were recently published and not included in the previous version of HRGM.

To construct HRGM2 as a human gut microbiome catalog of rigorous quality, we exclusively chose near-complete (NC) genomes. Initially, we identified genome chimerism using GUNC v1.0.1^7^. Genomes with clade separation scores (CSS) greater than 0.45 were excluded as chimeric genomes, where 0.45 serves as the default threshold for GUNC. Subsequently, we calculated the completeness and contamination of the genomes using a lineage-specific workflow of CheckM v1.0.18^13^. We selected genomes with completeness ≥ 90% and contamination ≤ 5% as NC genomes. To identify completeness-underestimated clades, we performed taxonomic annotation for all genomes using GTDB-Tk v1.7.0^6^ (database release GTDB R202^76^). Next, we define a clade as a completeness-underestimated clade if more than 95% of genomes within the clade, from phylum to genus, exhibit a completeness below 90%. Genomes within such clades were re-estimated for completeness and contamination by applying either bacterial or archaeal marker set, depending on their respective domain, to the taxonomic-specific workflow of CheckM. Genomes belonging to the phylum Patescibacteria were subject to quality reassessment using the CPR marker set^16^. Genomes meeting the criteria of completeness ≥ 90% and contamination ≤ 5% after reassessment were also considered NC genomes. The quality of these added genomes was validated with CheckM2 v0.1.2^15^.

### Species clustering and genome dereplication

We clustered NC genomes at the species level. Calculating pairwise distances for all genomes is excessively time-consuming, so we initially conducted preliminary clustering. For genomes with the same order according to GTDB taxonomic annotation, pairwise mash distances were calculated using Mash v2.3^77^ with the "-s 10000" parameter. We performed average-linkage-based hierarchical clustering with a 0.1^3^ cutoff to establish preliminary clusters. While the Mash algorithm is known for its speed, it may suffer from accuracy limitations when dealing with genome pairs exhibiting low coverage^78^. Therefore, we calculated average nucleotide identity (ANI) using ANImf^78^ for each pair of genomes within each preliminary cluster. To address potential ANI overestimation stemming from local alignment, we imposed a minimum coverage threshold of 0.6^79^ for each pair. If the coverage of a genome pair was below 0.6, we considered them to be different genomes. We then conducted average-linkage-based hierarchical clustering with an ANI threshold of 95% to cluster genomes at the species level^2,3,80^. To designate a representative genome within each species cluster, we computed a genome intactness score (*S*)^2,3^, *S = Completeness* - 5 x *Contamination* + 0.5 x log_lO_(*N*50), for each genome and selected the one with the highest *S* value as the representative. To eliminate duplicate genomes, we performed average-linkage-based hierarchical clustering within each species cluster using ANI 99.9% and coverage 0.81 as thresholds. The coverage threshold of 0.81 was calculated by considering the minimum completeness of the NC genome, which is 90%. This threshold is the product of the minimum completeness of the two genomes.

### Taxonomic annotation of species and construction of the phylogenetic tree

We performed taxonomic annotation of each species using GTDB-Tk v1.7.0^6^ (database release GTDB R202^76^) for a representative genome of each species. GTDB-Tk predicted 120 and 122 universal marker genes for bacterial and archaeal genomes, respectively, and conducted multiple sequence alignments (MSA) for concatenated sequences of the predicted marker genes of the representative genome for each species. Based on the MSA, we constructed a phylogenetic tree using IQ-TREE v2.2.0.3^81^. The phylogenetic tree was visualized using iTOL v6^82^.

### High-quality (HQ) genome annotation

We annotated the NC genomes as HQ if they contained 5S, 16S, and 23S rRNAs, as well as tRNAs for at least 18 out of the 20 possible amino acids. The 5S, 16S, and 23S rRNAs for each genome were predicted using Barrnap v0.9^83^. The parameters "--kingdom bac" and "-- kingdom arc" were applied for bacterial and archaeal genomes, respectively. An e-value threshold of 1e-05 was used for the prediction of 16S and 23S rRNAs, while an e-value threshold of 1e-04 was used for the prediction of 5S rRNA, considering its shorter length. For the tRNA search, tRNAscan-SE v2.0.9^84^ was used, with the "-B" option applied for bacterial genomes and the "-A" option for archaeal genomes.

### Identifying NC genome species overlap between HRGM2 and other catalogs

We investigated the NC genome species (Species containing NC genomes) overlap between HRGM2 and other catalogs using NC representative genomes. A species pair was deemed identical if the ANI > 95% and coverage > 0.6 between the representative genome of HRGM2 and that of another catalog. Due to the computational burden associated with pairwise comparisons for all representative genomes, we only compared representative genomes that have the same order according to the GTDB taxonomic annotation.

### Construction of pan-genomes for HRGM2

For 2,639 of the 4,824 species that contain more than one non-redundant genome, we constructed pan-genomes using Panaroo v1.3.0^85^ with the parameters "--clean-mode strict –c 0.90 -f 0.5 --merge_paralogs --core_threshold 0.90". The core genes were defined as present in at least 90% of genomes, otherwise, accessory genes. We used GFF files generated by Prokka v1.14.6^86^ as input to Panaroo. For each genome, Prokka was executed with the parameter "--kingdom Bacteria" or "--kingdom Archaea" depending on the domain.

### Construction of a species-specific marker gene database for HRGM2

The species-specific marker gene database for HRGM2 was constructed by following the methodology of MetaPhlAn4^8^. Candidates for species-specific marker genes were identified through the clustering method. Initially, gene sequences were collected from the pan-genomes of 2,639 species with multiple conspecific genomes. For the remaining species with one genome, all protein sequences from that genome predicted by Prokka were collected. The collected protein sequences were clustered into families with 90% identity using the "linclust^87^" function of MMseqs2 v13.45111^88^. Only protein families with lengths between 150 and 1,500 amino acids and averaging fewer than 1.5 copies per genome were included in subsequent steps. We identified core protein families for each species. "Coreness" was defined as the proportion of conspecific genomes within a species that possess a given protein family. A coreness threshold of 60% was set for species with fewer than 100 conspecific genomes, and 50% otherwise. Unique core protein families for each species were then identified. Core protein families that exhibited more than 1% coreness in other species were excluded from consideration. Finally, species-specific marker genes were identified from these candidates by alignment. The DNA sequences of these candidate genes were fragmented into 150 bp segments, which were aligned against all 155,211 non-redundant genomes using Bowtie2 with the "--sensitive" parameter. A candidate gene was considered to be present in a genome if at least one segment aligned to it. Based on this alignment, the coreness and uniqueness of the candidates were re-evaluated using the same criteria, and those still meeting these criteria were selected as species-specific markers. Up to 200 markers per species, which were the least shared with other species and the longest, were selected. Using these optimal markers, a custom database for MetaPhlAn4 was constructed using Bowtie2-build with the "--large-index" parameter.

### Simulation dataset generation for comparing taxonomic profiling methods

To compare taxonomic profiling methods based on HRGM2, we generated 5 Gbp simulation samples using CAMISIM v1.3^89^ with default parameters. Non-redundant genomes from HRGM2 were randomly selected to construct the simulation dataset. The complexity of the simulation community was systematically increased by selecting genomes in quantities of 40, 150, 600, 1,000, and 1,500. Additionally, the simulation data represented each species either as a single strain or as multiple strains. In the single strain scenario, one genome per species was chosen, whereas in the multiple strain scenario, up to five genomes per species were selected. Consequently, 10 different setups were organized based on the combination of five levels of community complexity and two strain strategies. Four samples were simulated for each setup, resulting in a total of 40 samples.

### Performance comparison of taxonomic profiling methods

We compared the performance of taxonomic profiling methods based on HRGM2 using simulation data. For marker-based taxonomic profiling, MetaPhlAn4 v4.0.3 was employed with the HRGM2 custom marker database. For DNA-based taxonomic profiling, Kraken2 v2.1.2^21^ and Bracken v2.8^22^ were utilized with the HRGM2 custom database. The HRGM2 Kraken2 custom database was constructed in two versions: "Representative" and "Concatenated”. The former was built with only the representative genome of each species, while the latter was constructed based on genomes concatenated with non-redundant genomes of each species. Kraken2 was run both with and without a confidence score of 0.2. For DNA-based taxonomic profiling, normalization of species abundance according to genome size was also considered. In the "Representative" version, species read counts were normalized by the representative genome size, whereas in the "Concatenated" version, by the average genome size of the species.

### Identification of subspecies-level geographic stratification via SNV-based analysis

To compare the number of detected single nucleotide variations (SNVs) between HRGM2 and UHGG, we first constructed reference SNV databases using non-redundant NC genomes from species represented by more than 10 genomes in both catalogs. SNV calling was performed with the genomes command in Maast v1.0.8^26^, using a minimum prevalence threshold of 0.9 and a minor allele frequency (MAF) cutoff of 0.01.

To investigate strain-level heterogeneity associated with geographic origin, we collated a total of 1,787 publicly available human fecal metagenome samples from 12 datasets (**Supplementary Table 6**), including 1,169 samples from Asia (India, Japan, and China) and 618 from Europe/US (Austria, Italy, Germany, France, and the United States). SNV-based analysis was conducted for species with more than 50 non-redundant NC genomes in HRGM2. Reference SNV databases were constructed using the workflow described above. Subsequently, SNV genotyping across all samples was performed using GT-Pro v1.0.1^27^. Pairwise genetic distances between strains were calculated, and phylogenetic trees were constructed by concatenating genotyped SNVs using the tree command in Maast. PERMANOVA test was applied to the distance matrices to assess subspecies-level separation between Europe/US and Asia. Species exhibiting significant geographic stratification were defined by a pseudo-F statistic > 30 and a PERMANOVA *p*-value < 0.01.

### Cataloging human gut microbial proteins and functional annotations

To construct a protein database for the human gut microbiome, we predicted coding sequences from NC genomes using Prokka. Proteins with identical sequences were pre-deduplicated, and clustering was performed at 100% identity using the "linclust" function of MMseqs2 with the "--min-seq-id 1" parameter, with other parameters set as previously described. Subsequently, clustering was sequentially repeated at 95%, 90%, 70%, and 50% identity by adjusting the "--min-seq-id" option to 0.95, 0.90, 0.70, and 0.50, respectively.

The representative protein sequences from the 90% identity clusters were functionally annotated using eggNOG-mapper v2.1.9^29^, based on the eggNOG protein database v5.0.2^90^. While eggNOG-mapper provides functional annotations for several terms, we employed additional annotation methods to enhance the overall annotation quality. First, we utilized DeepGOPlus v1.0.1^35^, which uses deep learning for gene ontology (GO) prediction, to annotate representative protein sequences from 90% identity clusters as in eggNOG-mapper. Among the results of GO predictions, only the predictions with a confidence score exceeding 0.3 were retained. Furthermore, we excluded the term "cellular anatomical entity (GO:0110165)", which refers to any structural component within a cell, as it annotated 89% of the protein clusters, leading to inflation of the prediction rate. Since DeepGOPlus was trained on data that includes protein sequences from eukaryotes, some predictions may correspond to functions specific to eukaryotic organisms, potentially leading to mispredictions for bacterial proteins. To address this, we retained only bacteria-associated GO terms in the predicted results. Using the UniRef90^91^ database downloaded in September 2023, we identified proteins derived from bacteria, and designated the GO terms associated with these proteins as bacteria-associated GO terms.

Second, we annotated carbohydrate-active enzyme (CAZyme) families using run_dbcan v4.1.4, the standalone tool of dbCAN3^37^, which employs a combination of HMMER, DIAMOND, and dbCAN_sub for prediction. We annotated 155,211 non-redundant genomes for the CAZyme families using the output of Prokka. To ensure higher confidence in the annotated results, only protein sequences predicted by at least two of the three approaches were classified into CAZyme families. Additionally, when the annotation results from each method differed, the results by HMMER were prioritized. If HMMER did not provide annotations, the results from dbCAN_sub were used instead.

Finally, we annotated defense functions against external threats, including antibiotics and viruses. Using the pan-genome of Panaroo outputs, we employed RGI v6.0.0^41^ with CARD v3.2.5 and WildCARD v4.0.0 to predict antibiotic resistance genes for 4,824 species. Through RGI, each gene family is annotated using the Antibiotic Resistance Ontology (ARO). Certain ARO categories encompass scenarios where multiple genes act in concert to confer antibiotic resistance. Typical examples include the glycopeptide resistance gene cluster (Van) and MexAB-OprM. In these cases, multiple genes work together to confer resistance. This cooperative relation between proteins was inferred by CARD ontology. To avoid overestimating the antibiotic resistance potential of a gene when only one component gene is present, we defined that all genes working together must be present for resistance to function. However, for glycopeptide resistance gene clusters, which are composed of regulatory, core, and accessory genes, the presence of antibiotic resistance is considered valid when all core genes are detected, even if accessory or regulatory genes are absent.

We analyze protein sequences from all 155,211 non-redundant genomes to predict the bacterial defense system against viruses using DefenseFinder v1.0.9^92^. We used a criterion of 80% to distinguish whether a defense system is present in the core or accessory genome for a species with more than 10 non-redundant genomes. Specifically, if more than 80% of the non-redundant genomes in a species have the system, it is classified as part of the core genome; otherwise, it is classified as part of the accessory genome.

### Comparison of GO prediction for human gut microbial proteins by DeepGOPlus and eggNOG-mapper

Representative protein sequences from the 90% identity clusters in the protein catalog were functionally annotated by GO term using eggNOG-mapper and DeepGOPlus. For comparison, the results of GO prediction by eggNOG-mapper were filtered by the same method as the DeepGOPlus, leaving only bacteria-associated GO terms. To determine which method resulted in more accurate predictions, we assessed how well the functional GO profiles of each species reflected its taxonomy. Functional GO profiles for each species were constructed from the predicted GO terms for representative genomes of each species by eggNOG-mapper or DeepGOPlus. Then, we used agglomerative clustering with Jaccard distance to cluster the GO profiles generated by each method. Agglomerative clustering requires a hyperparameter to determine the total number of clusters, so we increased this value from 1 to 1,000 to evaluate the homogeneity of the clustering results. We evaluated the performance by checking whether species belonging to the same taxonomic lineage—phylum, order, family, and genus—are grouped into the same cluster.

### Strain level comparison of CAZyme families between Western and non-Western countries

We first categorized countries in Europe and North America as Western and otherwise as non-Western. To identify species with different CAZyme families between strains from Western and non-Western countries, we analyzed profiles of the CAZyme family for all 155,211 non-redundant genomes. These profiles indicate the number of CAZyme families present in each genome. Using the PERMANOVA test with Euclidean distance, we found that genomes from Western countries had significantly different CAZyme profiles compared to genomes from non-Western countries within the same species. The "adonis2" function from the "vegan" package in R was used to identify species with significant differences in CAZyme profiles between Western and non-Western genomes.

### Assessing the importance of the NC genomes

To investigate the importance of NC genomes, we first identified 327 species that lacked NC genomes in UHGG but possessed NC genomes in HRGM2. For consistency, we considered representative genomes from each catalog as belonging to the same species if the ANI > 95% and the maximum coverage > 0.6. We then computed the non-overlapping genome proportion as 1 minus the coverage of each genome. To identify overlapping proteins, we used Prokka to predict proteins and clustered them using the "cluster" function of MMseqs2 with the option "--cov-mode 1 -c 0.8 --kmer-per-seq 80 --min-seq-id 0.50 --cluster-mode 2". The non-overlapping protein proportion was calculated by subtracting the overlapping protein proportion from 1.

To explore functions that are not primarily covered by the MQ genomes, we predicted KEGG modules^47^ for each representative genome of the 327 conspecific pairs using METABOLIC v4.0^93^. For KEGG modules detected in at least 100 species, we calculated the proportions observed exclusively in NC genome, solely in MQ genome, and in both.

To compare the functional composition of NC genomes and MQ genomes within species, we selected UHGG species with at least 10 non-redundant MQ genomes and NC genomes each. To reduce computational burden, we randomly selected up to 50 NC and 50 MQ genomes per species, focusing on species where the disparity in the number of NC and MQ genomes was less than twofold to minimize bias. As a result, we analyzed 543 species comprising 41,072 genomes. We downloaded the GFF files for these genomes, used Panaroo to identify gene clusters for each genome, and calculated the functional distance between genomes using the Jaccard distance metric.

### Assessment of metabolic independence

The metabolic independence of species was evaluated using 33 KEGG modules referenced in Watson et al^9^. For 4,824 representative genomes, KEGG modules were predicted using METABOLIC v4.0^93^. A representative genome was classified as a high metabolic independence (HMI) species if it possessed more than 80% of these 33 KEGG modules, and as a low metabolic independence (LMI) species if it possessed 20% or fewer. At the clade level, from phylum to genus, clades with 10 or more species where over half of the species were either HMI or LMI were designated as HMI clades or LMI clades, respectively.

### Reconstructing Genome-scale metabolic models and identifying metabolic interactions

The genome-scale metabolic models (GEMs) for each non-redundant genome were reconstructed using CarveMe v1.5.2^56^ with default parameters. On average, the reconstructed GEMs contained 889.30 metabolites, 1,267.77 reactions, and 810.54 gene-associated reactions. Due to the high computational burden of testing metabolic interactions for all possible species pairs in HRGM2 (11,633,076 pairs), we focused on predicting pairwise metabolic interactions for the top 1,000 species most prevalent in the human gut. To select these prevalent species, we utilized 1,160 metagenomic samples from healthy individuals, evenly distributed across various countries and collated for HRGM2 **(Supplementary Table 2)**. Based on our recommended taxonomic profiling method (Kraken2/Bracken with the “Concatenated” version database, confidence score of 0.2, and genome size normalization), we calculated relative abundances of species. A species was considered present in a sample if its relative abundance exceeded 1e-06, and its prevalence across all samples was determined.

For the top 1,000 species, the prevalence metrics were as follows: max = 100%, min = 28.53%, average = 60.34%, median = 56.25%. Pairwise metabolic interactions for these top 1,000 prevalent species were predicted using SMETANA v1.2.0^57^, with GEMs reconstructed from representative genomes and the parameters "--flavor bigg -g --molweight." Based on the distribution of cooperation scores (metabolic interaction potential, MIP) and competition scores (metabolic resource overlap, MRO) for 499,500 pairs, we identified cooperative and competitive pairs. Pairs were classified as cooperative if MIP > 3 and MRO < 0.667, and as competitive if MIP < 3 and MRO > 0.667. The mathematical formulas for MIP and MRO are described in the original SMETANA publication.

To identify metabolite driving cooperative interactions with the two low metabolic independence (LMI) orders, TANB77 and RF39, we reran SMETANA in detailed mode using the "-d" parameter (instead of "-g") on pairs that included either LMI order among the previously analyzed 499,500 species pairs. We focused on key metabolites essential for the survival of TANB77 and RF39, defined by a metabolite uptake score (MUS) > 0.9, which reflects the receiver’s growth dependency on a given metabolite. Extremely rare metabolites—those observed in fewer than 1% of pairs for each LMI order—were excluded from further analysis.

### Investigation of metabolic interactions among disease-associated species

We focused on microbial species enriched in Crohn’s disease (CD) and colorectal cancer (CRC), previously identified in our earlier study^64^. Disease-associated species were selected based on the following criteria: (1) Q value < 0.01, (2) |Log_2_ fold change| > 2, (3) Differential abundance (DA) direction = Disease (based on batch-corrected, integrated cohort), (4) inclusion among the top 20 species ranked by absolute average Shapley additive explanation (SHAP) values, and (5) SHAP direction = Disease.

Metabolic interactions among these species were examined using SMETANA from two perspectives: community-wise and pairwise. For the community-wise analysis, SMETANA was applied to the full CD- or CRC-associated species community and compared against 100 randomly generated communities of the same size, sampled from the top 1,000 prevalent species. In the pairwise analysis, all possible species pairs (N) within each disease-associated community were analyzed using SMETANA and compared to 2N randomly selected pairs from the same background set, providing increased statistical power.

To further characterize cooperative interactions in the CD-associated community, SMETANA was run in detailed mode. Following Machado et al^94^., SMETANA scores were aggregated across species pairs with identical donor-receiver configurations to represent overall influence on viability. Interactions were visualized in Cytoscape v3.10.3^95^. Key metabolites essential for receiver survival were identified based on MUS > 0.9. To validate whether these interactions manifest in actual CD samples, we used CD discovery cohorts from our previous study^64^. Due to small sample size, batch correction was applied via MMUPHin v1.16.0^96^, and all CD samples were pooled. Samples were stratified by the median relative abundance of key donor species.

For CRC, competitive interactions were explored using each species’ individual media, representing its nutritional requirements, extracted via SMETANA’s mro_score function. To assess how competition impacts species dominance in the community, we analyzed CRC discovery cohorts^64^, calculating the relative abundance of the 11 CRC-associated species per sample. A species was considered present if its relative abundance exceeded 1e−6 among all HRGM2 species. Unlike CD, the larger sample sizes in CRC cohorts allowed for stratified analysis by cohort.

## Data availability

By accessing the web server, www.decodebiome.org/HRGM2/, users can browse and download all genomes for representative species, their annotations, and metadata, including geographical origin, taxonomy, genomic content, and genome statistics. The five classes of protein catalogs, 16S rRNA sequences, and SNVs are also provided with their functional annotation and taxonomic origin. In addition to publicly available datasets, we incorporated three newly generated datasets (PRJNA1227720, PRJNA1227423, and PRJNA1226738), with full details in **Supplementary Table 1** and raw metagenomic sequencing data available in the NCBI Sequence Read Archive.

## Code availability

The source code utilized for the construction and analysis of HRGM2 is publicly available at https://github.com/netbiolab/HRGM2.

## Extended Data Figure legends

**Extended Data Fig. 1.**
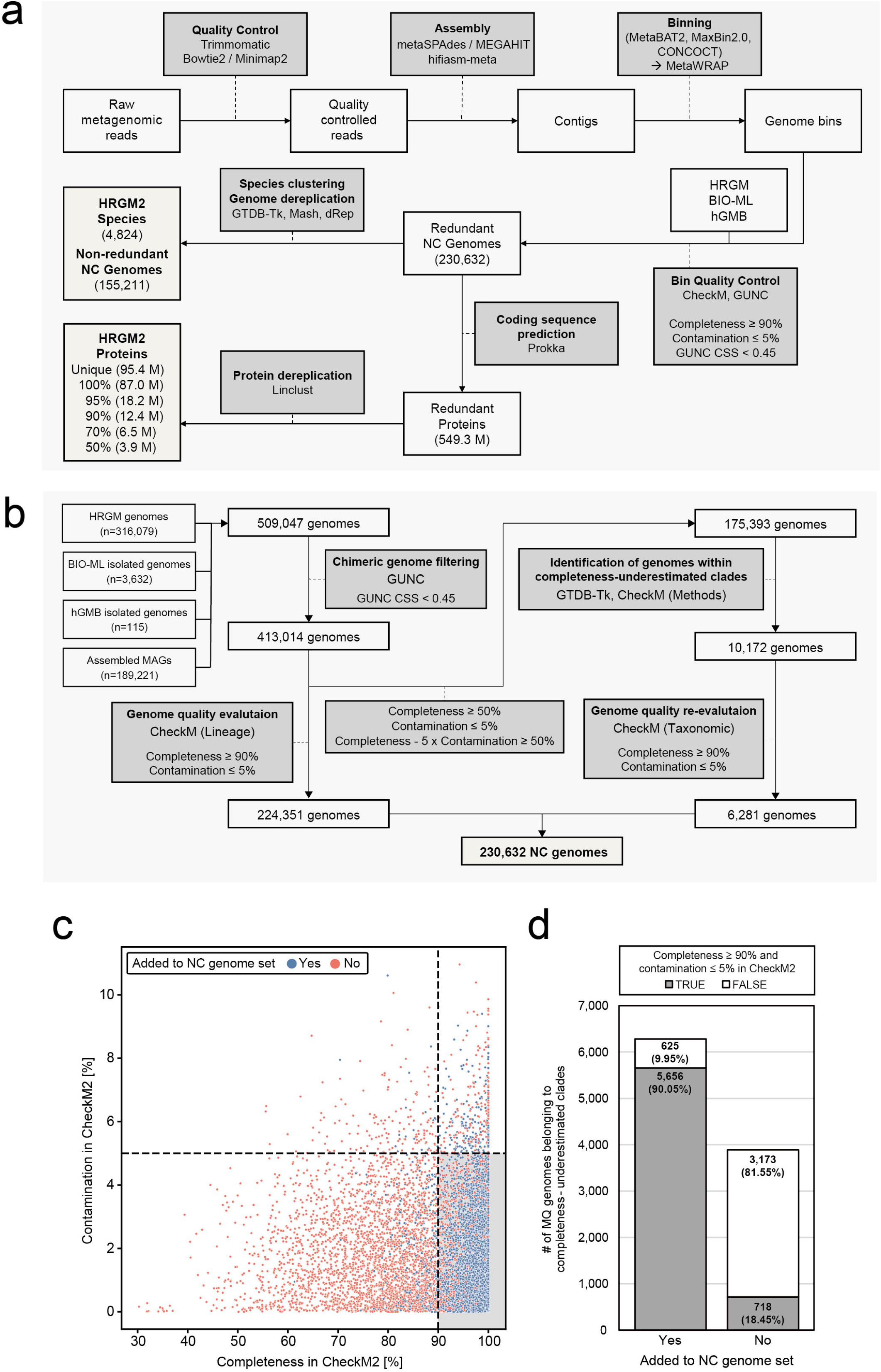
Bioinformatics pipelines for HRGM2 construction. **a**, The overall pipeline for HRGM2 construction. **b**, Pipeline to control genome quality. **c**, CheckM2 assessment of 10,172 MQ genomes previously assessed by CheckM (completeness-underestimated). The 6,281 genomes that were added to the near-complete (NC) genome set based on assessment using universal bacterial markers and CPR markers are marked with blue dots. The vertical and horizontal dashed lines indicate the 90% completeness and 5% contamination threshold, respectively. The gray area includes genomes that meet the NC genome criteria according to CheckM2. **d**, Count and proportion of filtered in and out by universal bacterial markers and CPR markers that meet the NC criteria under CheckM2.

**Extended Data Fig. 2.**
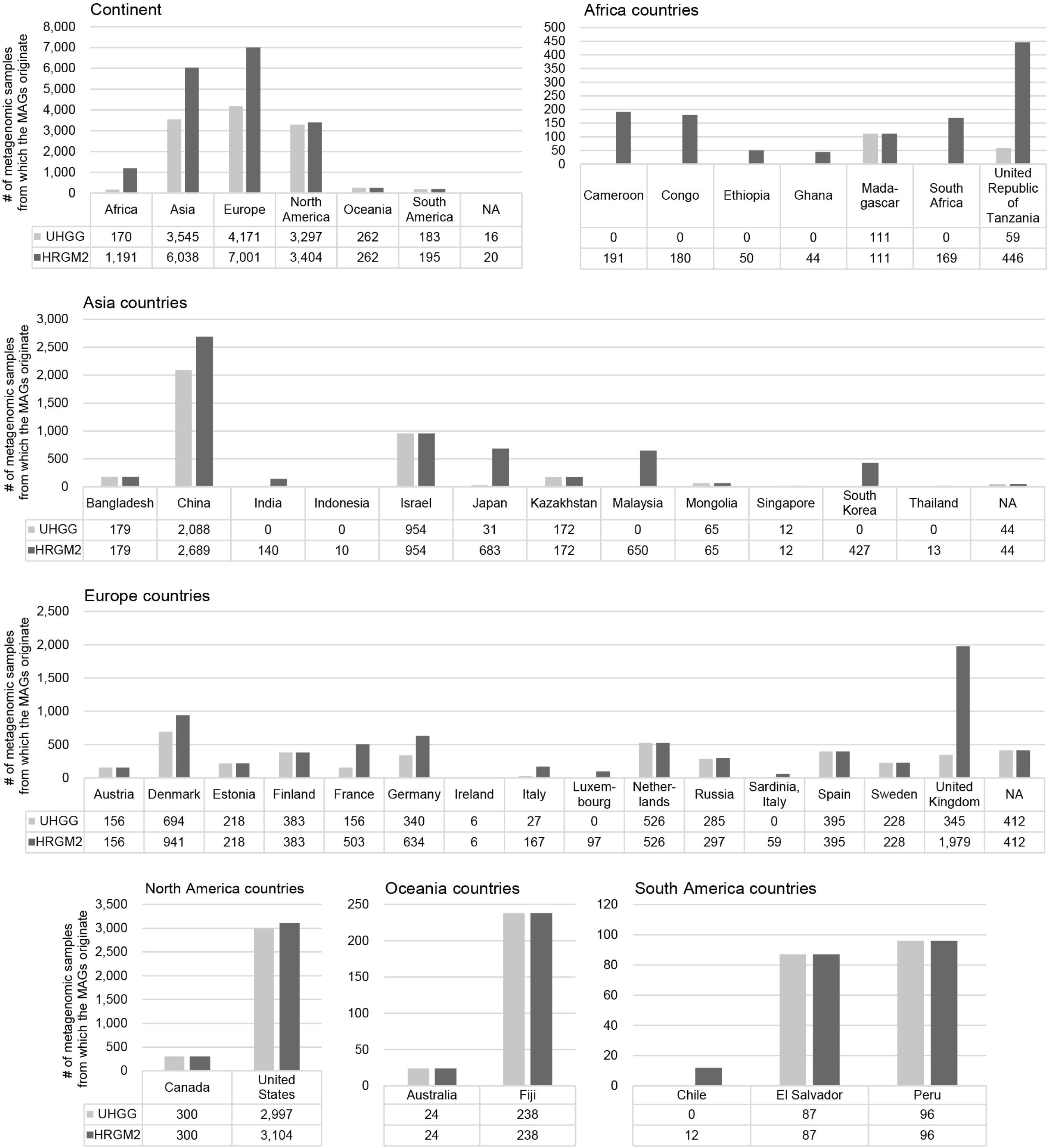
The number of metagenomic samples from which the MAGs originate in UHGG and HRGM2 by continent and country.

**Extended Data Fig. 3.**
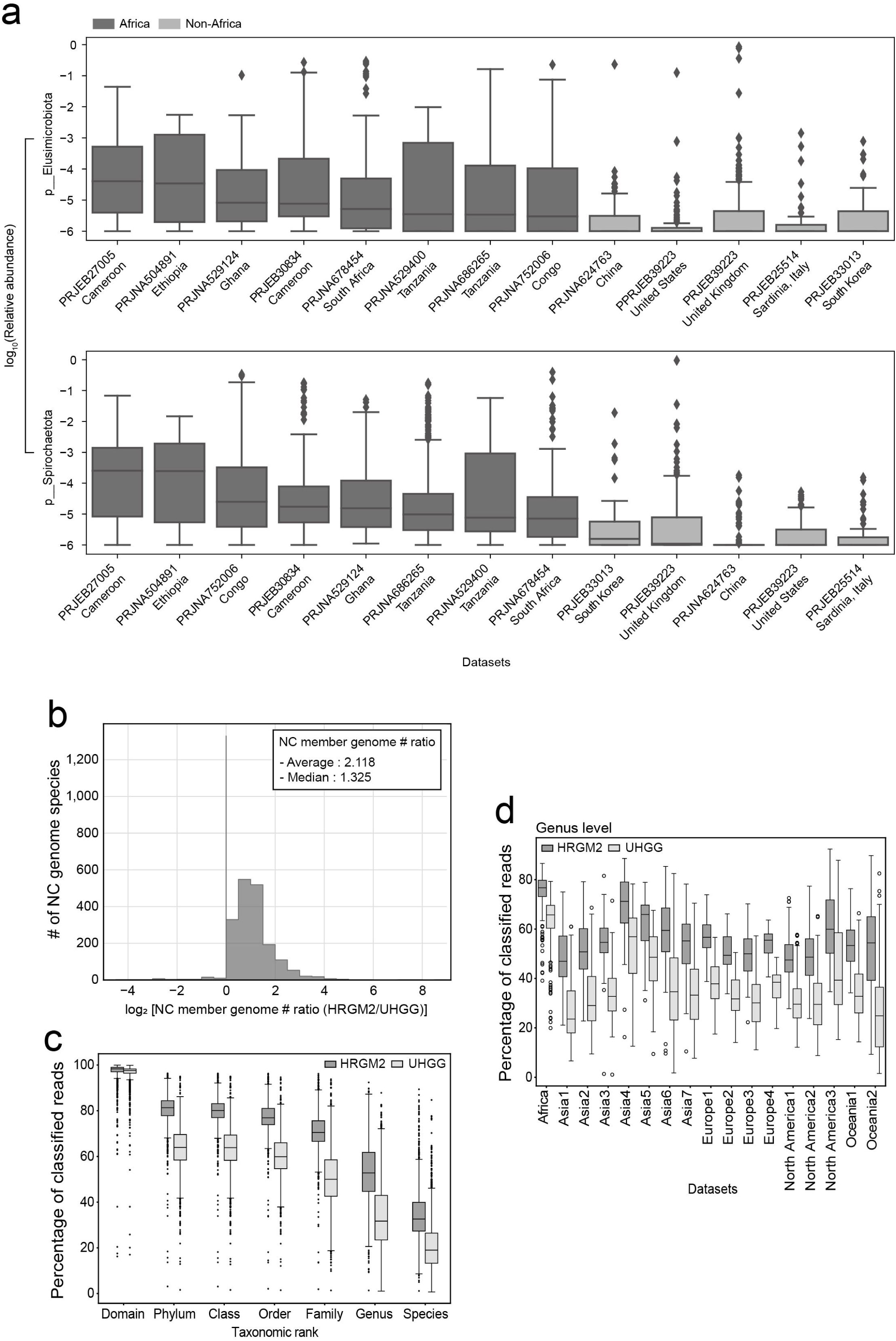
Extended comparative analyses supporting the HRGM2 overview. **a**, Abundance distributions of phyla Elusimicrobiota (up) and Spirochaetota (down) across Africa and non-Africa datasets. All African datasets used for the construction of HRGM2 are included. For comparison, five non-African datasets were randomly selected. The datasets are represented by study accession and country. In the case of “PRJEB39223, United Kingdom”, 400 samples were randomly selected due to the large total sample size. Relative abundances were batch-corrected using MUPPHin and log_10_-transformed. A phylum was considered present in a sample only if its relative abundance exceeded 1e-06; values below this threshold were replaced with 1e-06. Thus, a y-axis value of -6 indicates absence of the phylum in that sample. Boxes were sorted from left to right, in descending order by median. **b**, Comparison between HRGM2 and UHGG for the number of NC member genomes for conspecific pairs. **c-d**, Comparison of the percentage of classified reads between HRGM2 and UHGG at (c) each taxonomic rank and (d) the genus rank, with datasets stratified by continent. For boxplots, box lengths represent the interquartile range of the data, and whiskers extend to the lowest and highest values within 1.5 times the interquartile range from the first and third quartiles, respectively. The center bar represents the median. All the outliers are shown in the plots.

**Extended Data Fig. 4.**
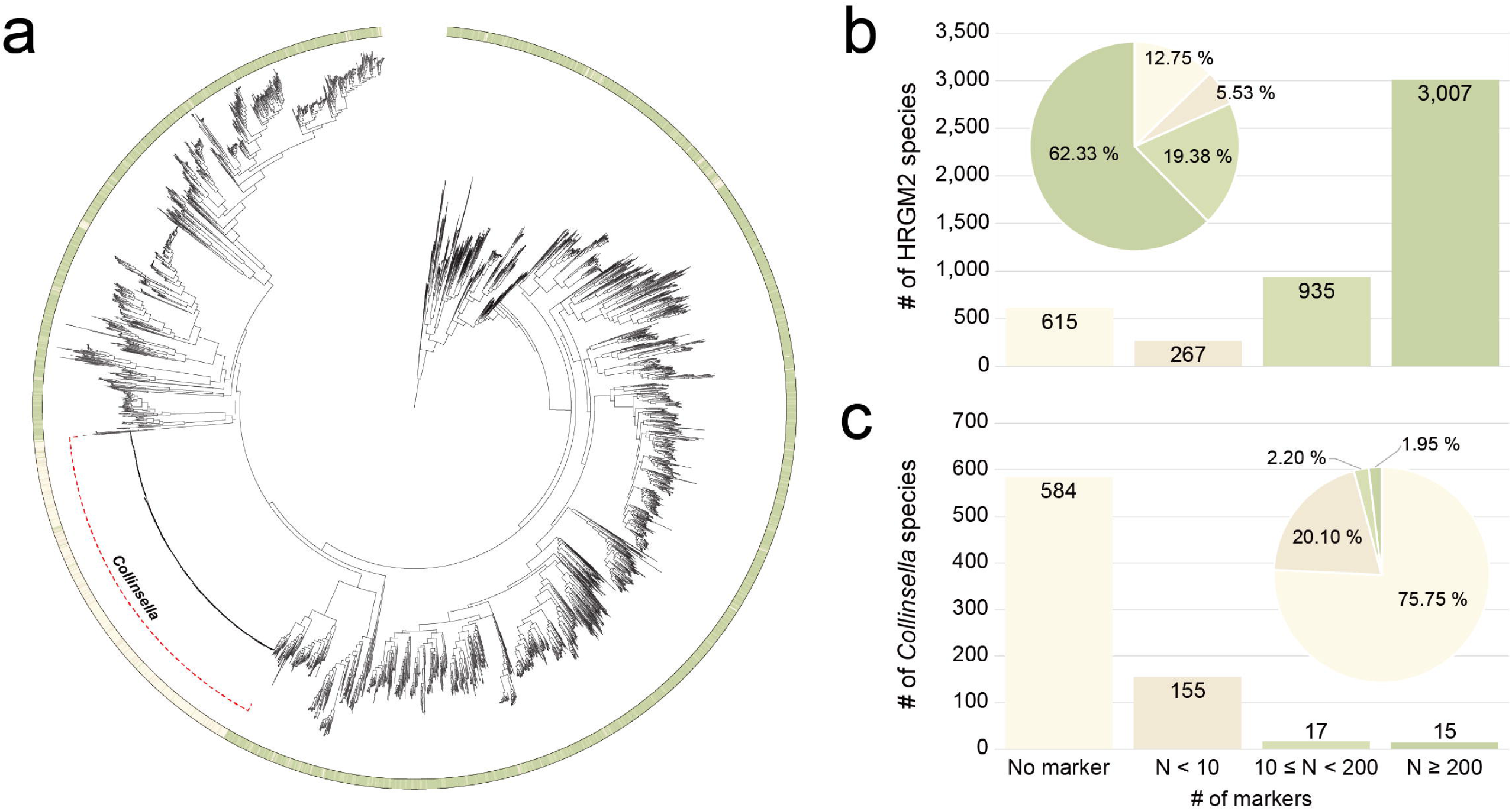
Summary of HRGM2 species-specific marker gene database. **a**, Maximum-likelihood phylogenetic tree with annotations of the number of species-specific marker genes. The color of the strip according to the number of species-specific marker genes is the same as in (b, c). **b**-**c**, The number and proportion of species in (b) HRGM2 and (c) *Collinsella* genus according to the number of species-specific marker genes.

**Extended Data Fig. 5.**
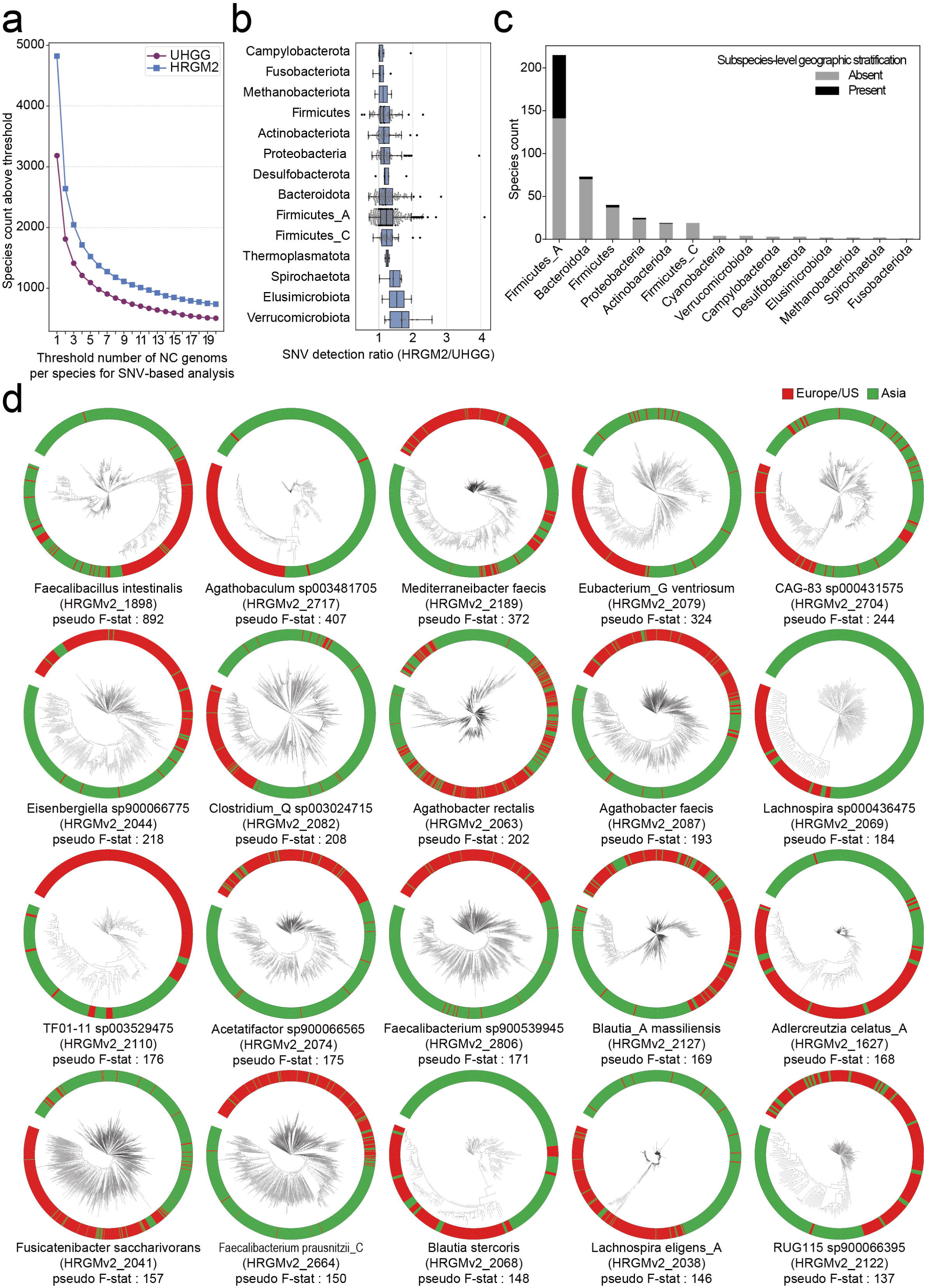
Analysis of strain-level heterogeneity across geographic regions in the human gut microbiota. **a**, Number of species exceeding varying thresholds of near-complete (NC) genomes required for SNV-based analysis. HRGM2 consistently retains more species than UHGG across all thresholds, demonstrating enhanced capacity to resolve subspecies-level genetic heterogeneity. **b**, Comparison of the number of SNVs identified in species with ≥10 non-redundant NC genomes present in both catalogs. Boxplots show the distribution of SNV detection ratios (HRGM2/UHGG) per species. Box lengths represent the interquartile range of the data, and whiskers extend to the lowest and highest values within 1.5 times the interquartile range from the first and third quartiles, respectively. The center bar represents the median. All the outliers are shown in the plot. **c**, Number of species within each phylum exhibiting subspecies-level geographic stratification between Europe/US and Asia (black bars), based on PERMANOVA test (*p-value* < 0.01 and pseudo-F statistic > 30). **d**, Representative phylogenetic trees constructed from SNVs identified in metagenomic samples for top 20 species with geographic stratification, colored by geographic origin (green: Asia; red: Europe/US). Species names, HRGM2 species identifiers, and the corresponding pseudo-F statistics are shown below each tree.

**Extended Data Fig. 6.**
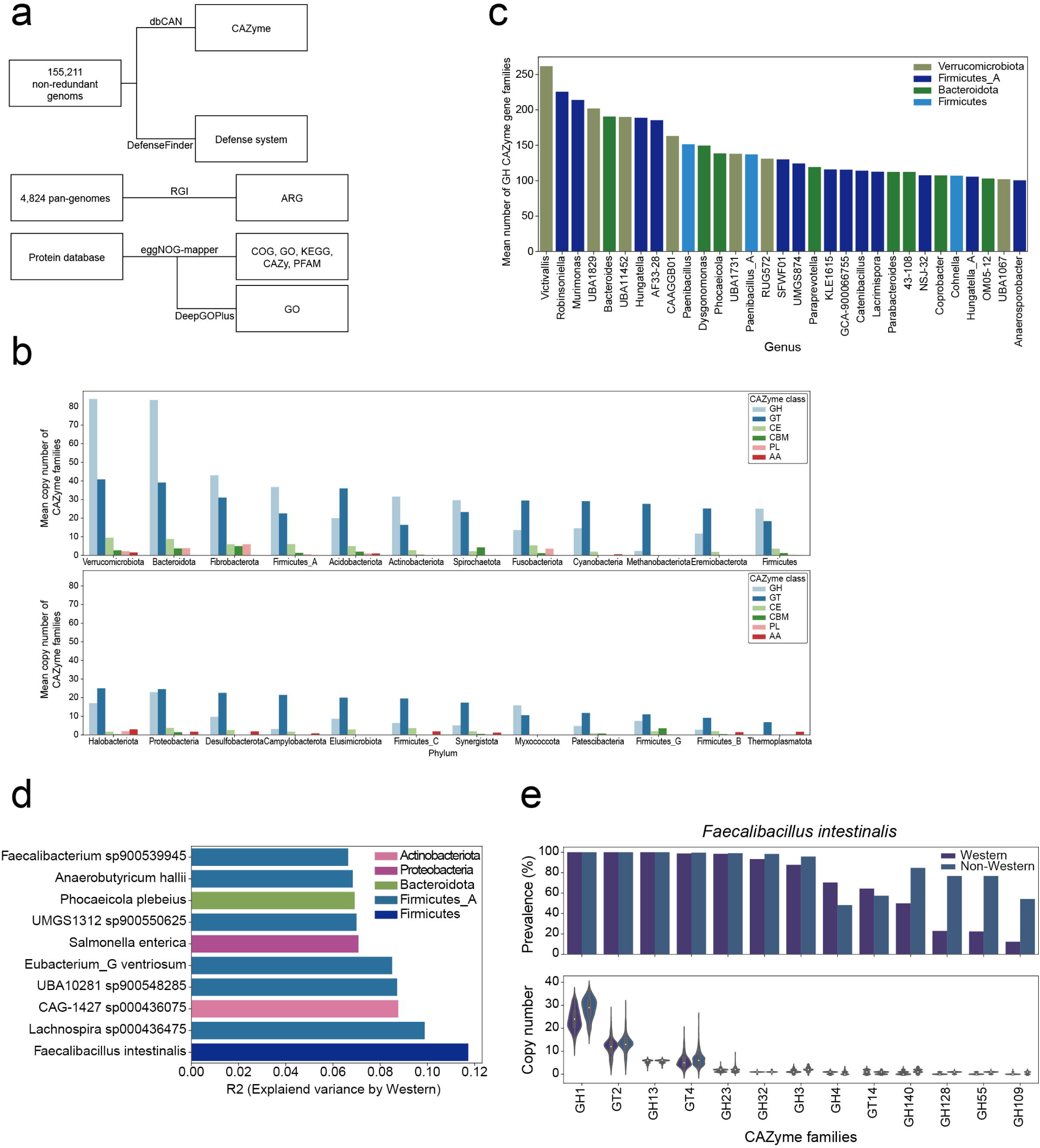
Functional landscape of human gut microbiome. **a**, Summary of functional prediction pipeline in HRGM2. **b**, Average copy number of CAZyme families per phylum. GH: Glycoside Hydrolase, GT: Glycosyl Transferase, CE: Carbohydrate Esterase, CBM: Carbohydrate Binding Module, PL: Polysaccharide Lyase, AA: Auxiliary Activity. **c**, Average copy number of GH CAZyme families for genera with more than 100 GH CAZyme families. **d**, Explained variance by Western/non-Western categorization for the 10 species with the most distinct CAZyme profiles between Western and non-Western continents. In (**c**) and (**d**), the color of the bars represents phylum. **e**, Comparison of the prevalence and the copy number of each CAZyme family between *Faecalibacillus intestinalis* genomes from Western and non-Western countries.

**Extended Data Fig. 7.**
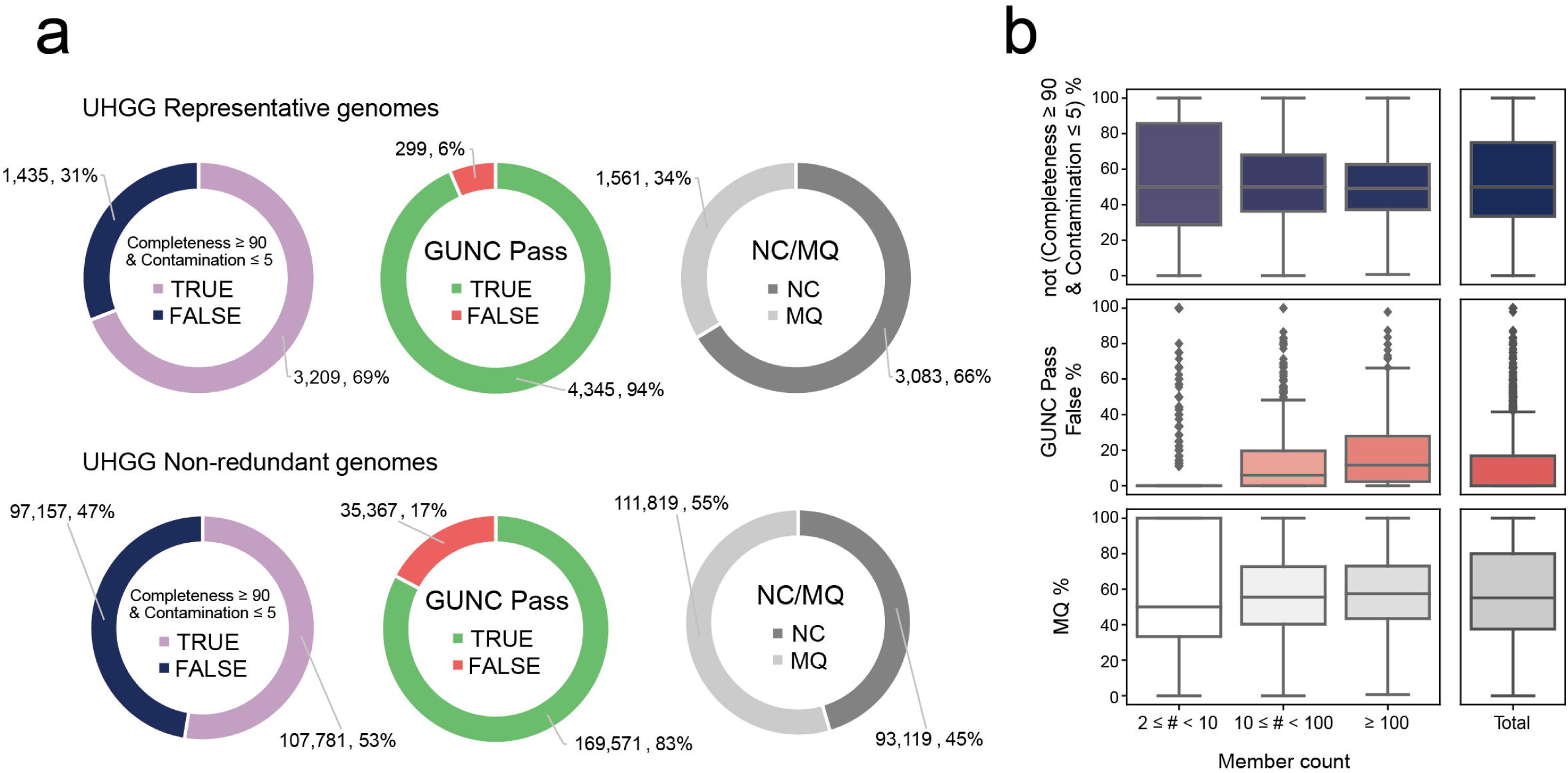
Summary of genome qualities for UHGG. **a**, Number and proportion of genomes with completeness ≥ 90% and contamination ≤ 5% (left pie chart), that passed GUNC (center pie chart), and that met the NC criteria (right pie chart), in UHGG representative genomes (up) and non-redundant genomes (down). **b**, Distribution of the percentage of genomes that are not completeness ≥ 90% and contamination ≤ 5% (top), that did not pass GUNC (middle), and that did not meet the NC criteria (bottom) for each UHGG species with at least two non-redundant genomes. The distributions are either categorized by the number of non-redundant genomes included in each species (left) or not (right) (2 ≤ # < 10, n = 1,590; 10 ≤ # < 100, n = 877; ≥ 100, n = 319; Total, n = 2,786). Box lengths represent the interquartile range of the data, and whiskers extend to the lowest and highest values within 1.5 times the interquartile range from the first and third quartiles, respectively. The center bar represents the median. All the outliers are shown in the plots.

**Extended Data Fig. 8.**
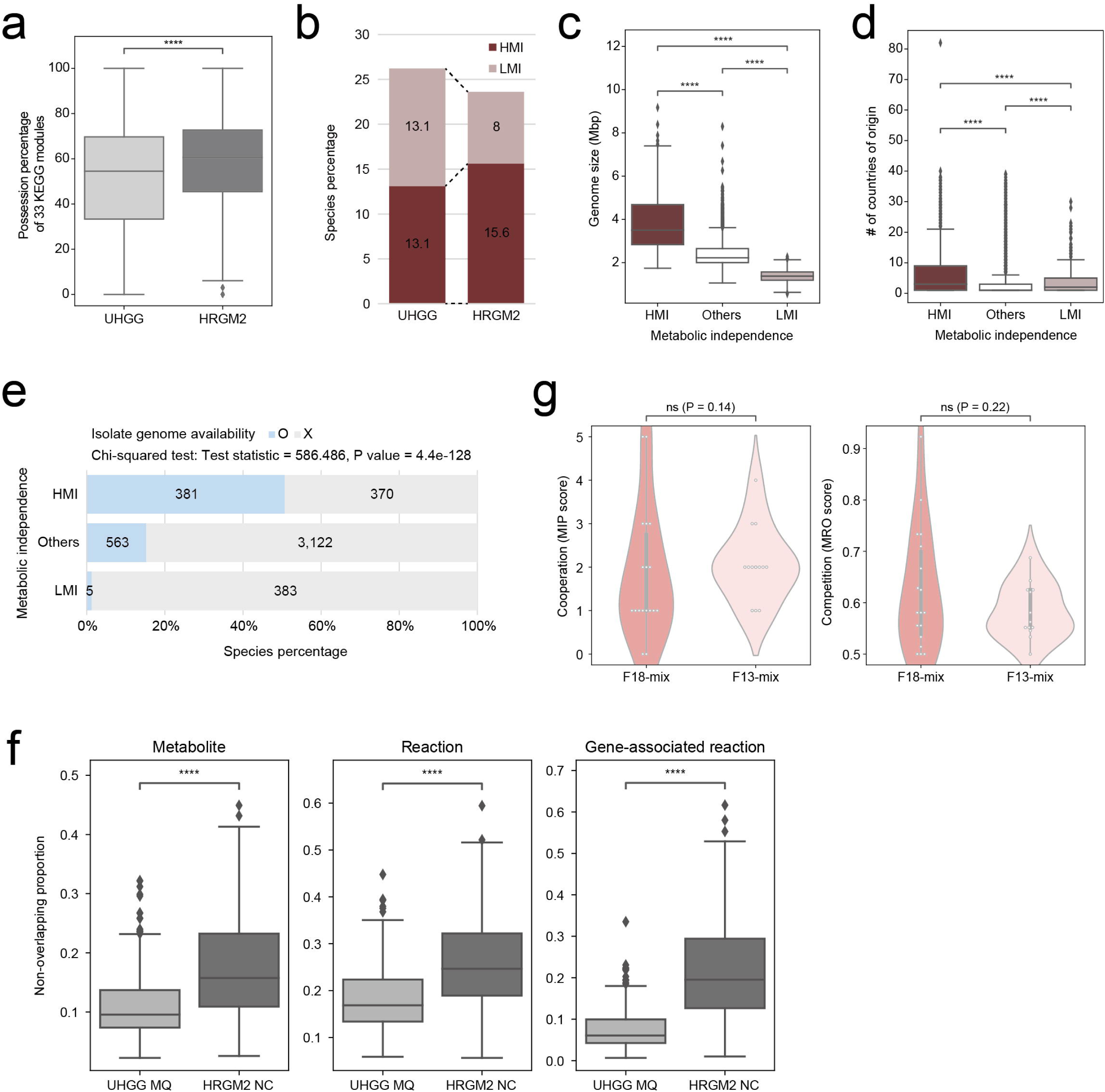
Additional analyses for metabolic independence and interaction. **a**, Possession percentage of 33 KEGG modules for UHGG (n = 4,644) and HRGM2 (n = 4,824) species. Statistical significance was assessed using a two-sided Mann-Whitney U test (P = 1.74e-42), indicating notable differences in module possession between the two catalogs. **b**, HMI and LMI species percentage in UHGG and HRGM2. **c**, **d**, Representative genome size (**c**) and the number of countries where each species originated (**d**) of HMI, other, LMI species. In (c) and (d), the number of HMI/Others/LMI species is 751/3,685/388 and 688/3,533/383, respectively; (d) considers only species with country information. *P*-value was calculated using a two-sided Mann-Whitney U test (P for HMI-Others/Others-LMI/HMI-LMI = (c) 3.270e-227/1.198e-205/2.297e-168; (d) 6.308e-29/1.839e-05/3.472e-05). **e**, Number and percentage of species with available isolate genomes, categorized by metabolic independence. The association between metabolic independence and the availability of isolate genomes was assessed by a Chi-squared test (two-sided). **f**, Comparison of the proportions of non-overlapping metabolites (left), reactions (middle), and gene-associated reactions (right) between 327 conspecific MQ and NC GEMs. Differences were evaluated with two-sided Wilcoxon signed-rank test (P for Metabolite/Reaction/Gene-associated reaction = 3.078e-19/1.203e-26/9.691e-45). **g**, Violin plot showing distribution of MIP and MRO scores for F18-mix (n = 18) and F13-mix (n = 13) strains of Kp-2H7. *P*-value was calculated using a one-sided Mann-Whitney U test. For boxplots in Extended Data Fig. 8, box lengths represent the interquartile range of the data, and whiskers extend to the lowest and highest values within 1.5 times the interquartile range from the first and third quartiles, respectively. The center bar represents the median. All the outliers are shown in the plots. ****, P < 1e-04; ns: not significant, P > 0.05.

